# A transitional senescence program drives inflammatory monocyte state expansion in aging humans

**DOI:** 10.64898/2026.09.09.750459

**Authors:** Themistoklis Vasilopoulos, Paolo S. Turano, Luis Garza-Martínez, Elizabeth Akbulut, Mounika Konda, Utz Herbig, Patricia Fitzgerald-Bocarsly, Ricardo Iván Martínez-Zamudio

## Abstract

Dysfunctional monocyte states contribute to age-related pathologies and systemic inflammation. However, the gene regulatory networks governing the transition to these states remain unknown. Here we used bulk and single-cell multidimensional integrative profiling to reveal previously uncharacterized monocyte state transitions during human aging. We show that a transient senescent-like population arising from classical CD14^++^ CD16^-^ monocytes drives the accumulation of an inflammatory monocyte state in aging humans. This senescence-associated transition is orchestrated by the master senescence regulator AP-1, which acts on a pre-established chromatin landscape to rewire the monocyte transcription factor (TF) network and activate both senescence– and age-associated inflammatory transcriptional programs. Through integration with clinical transcriptomic datasets, we demonstrate that senescent-like and aged monocytes are transcriptionally primed toward sepsis-associated states. Overall, our study provides the core gene-regulatory principles underlying a senescent-like transitional state in monocytes and identifies AP-1 as an attractive target to modulate systemic inflammation in age and disease.

## Introduction

Aging, the major driver of chronic disease, morbidity and mortality, is defined by the gradual loss of tissue function and characterized by low level chronic inflammation^1,2^. Critical to the aging process is the progressive transition of cells across multiple tissues towards a senescent phenotype characterized by a stable proliferative arrest and a senescence-associated secretory phenotype (SASP) that generates an inflammatory microenvironment^3,4^. Although senescent cells are immunogenic and are physiologically removed from tissues by the innate and adaptive immune systems^5^, accumulating evidence shows that the cellular components of the immune system become progressively dysfunctional with age^6^. This process, known as immunosenescence is characterized by a marked decline in plasmacytoid dendritic cell (pDCs) function and numbers^7^, increased myeloid-biased hematopoiesis as well as chronic inflammation^8,9^. While recent efforts have focused on characterizing dysfunctional states, such as senescence^10–13^ and exhaustion^14,15^, within adaptive immune compartments, whether the innate immune system undergoes age-associated, senescence-like state transitions is poorly understood. Unraveling the molecular mechanisms that drive innate immune system dysfunction during aging is likely to reveal opportunities to restore immune cell function and extend healthspan in older populations.

Monocytes are a short-lived^16^, bone marrow-derived, circulating innate immune cell population that plays critical roles in immunosurveillance. Monocytes participate in the initial response to infection and tissue damageand coordinate adaptive immunity by virtue of their antigen-presentation capacity and their differentiation potential into macrophages or some dendritic cell lineages^17,18^. Three main circulating monocyte subsets with distinct functions have been characterized^19^: an abundant (80-85%) and highly migratory classical CD14^++^CD16^-^ population specialized in pathogen recognition and phagocytosis^20–23^, an intermediate CD14^++^CD16^+^ (5-10%) population that mediates robust inflammatory signaling output and antigen presentation^24–27^ and non-classical CD14^+^CD16^++^ monocytes (5-10%) that perform intravascular anti-viral surveillance functions^28^. Accumulating evidence shows age-associated dysregulation across the three monocytes populations^29^, including the expansion of the intermediate and non-classical subsets^30–32^, elevated basal secretion of inflammatory mediators^33,34^, altered surface display of chemokine receptors^35^, and impaired Toll-like receptor (TLR)-dependent signaling^36^, phagocytosis^33^ and metabolism^37,38^. This age-associated transition towards a dysfunctional, inflammatory state is reminiscent of senescent phenotypes. However, while recent high-throughput studies have established correlations between transcriptomic, DNA methylation and proteomic profiles with functional defects^34,39–41^, the gene-regulatory mechanisms that drive senescence– and inflammation-associated monocyte transitions during human aging are not known.

In this study, we combined senescent cell isolation with bulk and single-cell (sc) multiomic profiling of monocytes from younger and older individuals. This high-resolution approach uncovered a distinct, transitional population of monocytes with senescence features that fuels the expansion of an inflammatory monocyte state in aging humans. Mechanistically, we show that the master senescence regulator AP-1^42,43^ leverages a pre-established chromatin landscape to rewire transcription factor (TF) network connectivity to drive senescence– and age-associated inflammatory gene expression. Crucially, integration with clinical transcriptomic cohorts shows that the senescent as well as the inflammatory monocytes that accumulate in older individuals are transcriptionally skewed toward sepsis-associated states. Together, our findings define the gene-regulatory principles underlying the expansion of inflammatory monocyte populations during human aging and reveal promising inroads for the therapeutic management of monocyte-driven systemic inflammation.

## Results

### Age-associated increase in senescence-associated β-galactosidase activity CD14-positive monocytes of older humans

We analyzed a large dataset of peripheral mononuclear blood cells (PBMCs) isolated from young (n=66; ages 20-39) and older (n=180; ages 60-94) donors labeled with a fluorescent senescence-associated β-galactosidase (SA-βGal) substrate, as this approach proved effective in identifying PBMC subsets with senescence features^11^. After extracting the flow cytometric profiles of CD14-positive monocytes, including CD16-positive subpopulations (**Extended Data Figure 1a**), quantification of the log_10_-transformed median fluorescence intensity (MFI) did not show a significant change in the average SA-βGal activity in monocytes between age groups but rather a significant increase in its variance in older donors (**Figure 1a**), consistent with the age-associated increase in immune variation^44^. Because our initial analysis did not explicitly evaluate CD16 expression and CD16-positive monocyte subtypes have distinct functions^17^, we performed an additional experiment in PBMCs isolated from a cohort of 6 young and 6 older humans. We evaluated SA-βGal activity, CD14 and CD16 expression as well as mitochondrial membrane potential (MitoMP) and mass (MitoBright) and analyzed their signal intensity profiles by flow cytometry. After normalization, scaling and debatching of the data, quantification of CD14 and CD16 MFIs using a traditional gating approach showed increased variability in the distributions of all three monocyte populations (CD14^++^CD16^-^, CD14^++^CD16^+^ and CD14^+^CD16^++^) in the older group, although this was not statistically significant at the donor level (**Extended Data Figures 1b,c**). We then performed an unbiased single-cell analysis using the cyCONDOR platform, which simultaneously captures continuous marker signal distribution and clusters cells in high-dimensional space^45^. The signal intensity and distribution of CD14 and CD16 across monocytes were the main drivers of the uniform manifold approximation and projection (UMAP) (**Figure 1b**). Assessment of mitochondrial membrane potential and mass revealed a gradient of mitochondrial electrical output and a uniform mitochondrial size distribution that were not significantly affected by age (**Extended Data Figures 1d-f**), consistent with the metabolic heterogeneity of monocytes^46^ and ruling out gross mitochondrial defects developing as a function of age. In contrast, we observed a significant age-associated increase in SA-βGal activity in monocytes, mostly CD14^++^CD16^-^, across the UMAP (**Figures 1c,d**), although there were donor-specific differences (**Figure 1e**). Six clusters, defined largely by CD14 and CD16 signal intensities, exhibited age-specific proportion shifts that reflected an expansion of the CD14^++^CD16^+^ population in older donors (**Figures 1f,g**), in line with the results of manual gating quantification. Within these clusters, the SA-βGal activity increase in monocytes from older donors originated mostly from the CD14^++^CD16^-^ population (cluster 1) and a subpopulation of CD14^++^CD16^+^ monocytes (cluster 2) (**Figures 1h,i**). We obtained similar results in a second cohort of 8 young and 8 older donors (**Extended Data Figures 1g-k**). Based on this extensive flow cytometric high-dimensional analysis, we conclude that there is an age-associated increase of SA-βGal activity in CD14^++^CD16^-^ as well as in a subpopulation of CD14^++^CD16^+^ monocytes of older individuals.

**Figure 1.**
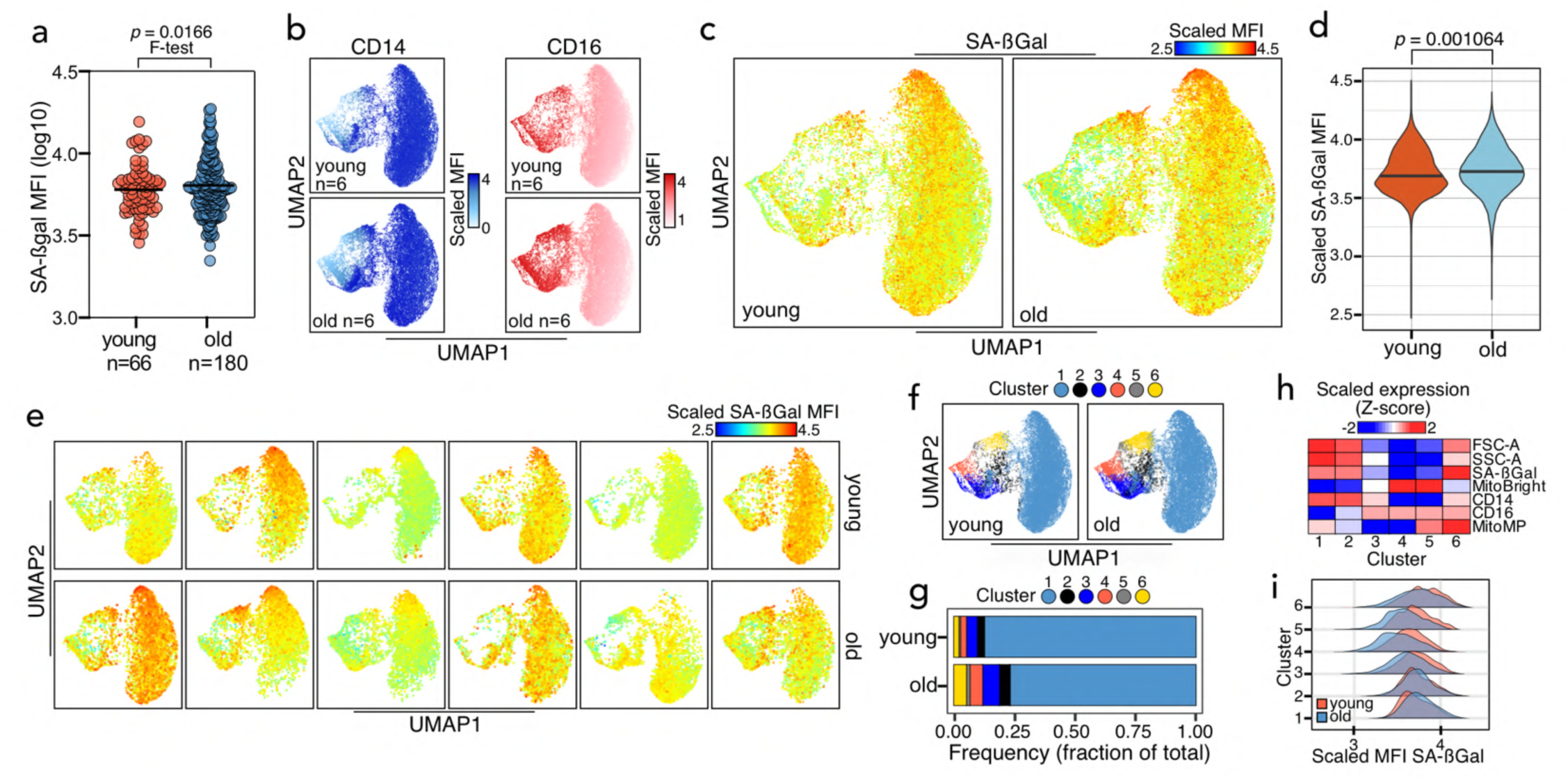
Increased SA-βGal activity in monocytes of older humans. **a.** Scatter plots of log10-transformed MFI values for SA-βGal activity in the CD14-positive monocyte compartment in the indicated number of younger and older donors. Each circle represents a donor and the black lines represent the mean. *P*-values were determined using a two-tailed unpaired Student’s *t*-test with Welch correction (*p* = 0.1611) and an F-test of variance equivalence (shown on top of panel). **b.** Uniform Manifold Approximation and Projections (UMAPs) showing the scaled MFI values for CD14 and CD16 from the indicated number of younger and older donors. **c-e.** Aggregate (c), individual (e) UMAPs for SA-βGal scaled MFI showing an increase of signal in older donors, and violin plot showing the distribution shifts of scaled SA-βGal MFI values between younger and older donors (e). *P*-value was determined using a Wilcoxon rank-sum test from a sample of 1,000 random cells per age group from a total dataset of 60,000 monocytes (5,000 cells per donor per age group). **f.** UMAP showing FlowSOM clusters determined from input variables FSC-A, SSC-A, CD14, CD16, SA-βGal, MitoMP and MitoBright. **g.** Frequency histogram showing the FlowSOM cluster distribution shifts between younger and older donors. **h.** Heatmap showing the Z-score expression of the FlowSOM input features per UMAP cluster across all samples. **i.** Density plots showing the shifts in distribution of scaled SA-βGal MFI values across FlowSOM clusters from f.

### Largely non-overlapping senescence and aging transcriptomes of CD14-positive monocytes

To further characterize monocytes with increased SA-βGal activity during human aging, we isolated CD14-positive monocytes with low, medium and high SA-βGal activities from 7 young and 7 older donors and subsequently profiled their bulk transcriptomes (**Extended Data Figure 2a**; see Methods for details). After controlling for donor variability and batch effects, unsupervised correlation, principal component (PC) and self-organizing map (SOM) analyses of total reads on exons revealed a clear separation and distinct expression portraits of SA-βGal-high relative to SA-βGal-low/-med CD14-positive monocytes (**Figures 2a-c**). We then performed differential expression analysis, identifying 902 SA-βGal activity-specific differentially expressed genes (DEGs). Weighted gene co-expression network analysis (WGCNA)^47^ of SA-βGal activity-specific DEGs generated two modules, blue and turquoise, specific to SA-βGal-low/-med and SA-βGal-high CD14-positive monocytes, respectively. The expression of DEG components in these modules exhibited variability in an SA-βGal activity-dependent manner and was largely independent of donor age (**Figures 2d,e**). Functional overrepresentation analysis of the SA-βGal-low-specific blue module revealed enrichment of pathways consistent with monocyte function, including IL2 and IL6 signaling pathways and coagulation^17^. In contrast, DEGs of the turquoise, SA-βGal-high-specific module, enriched for P53 pathway and IFNα/γ response, pathways that are overrepresented in the transcriptomes of senescent cells^10,48,49^ (**Figure 2f**). We also tested and found an upregulation at the protein level of a set of classical senescence markers 53BP1, CDKN1A (p21), CDKN1B (p27) and JUN, a member of the AP-1 family of master regulators of senescence transcriptional programs across multiple cell types^42,50–52^, in an SA-βGal activity-dependent manner in one young and one older donor (**Extended Data Figure 2b**). An equivalent analysis focusing on the effects of age on the monocyte transcriptome revealed age-specific expression portraits (**Extended Data Figure 2c-e**) and identified 740 age-specific DEGs distributed across two modules, I and II, whose expression patterns varied exclusively with age and independently of SA-βGal activity levels (**Extended Data Figures 2f,g**). DEGs in module I characterizing young monocytes enriched for proliferation-associated MYC and E2F targets, whereas module II DEGs that defined older monocytes enriched for several inflammatory and secretory pathways (**Extended Data Figure 2h**). Given the functional similarity of enriched pathways of SA-βGal-high and older CD14-positive monocytes, we reasoned that a large overlap in the respective DEGs should exist. Contrary to this expectation, there was little overlap between SA-βGal activity– and age-specific DEGs (**Figure 2g**). Plotting the raw counts of a subset of SA-βGal activity– and age-specific DEGs confirmed the cell state-specific gene expression patterns (**Figure 2h** and **Extended Data Figure 2i**). Taken together, our comprehensive transcriptome analysis demonstrates that senescence and aging transcriptomes of CD14-positive monocytes are largely non-overlapping and suggest the existence of two distinct senescent and aged CD14-positive monocyte populations and/or a monocyte state transition (see below).

**Figure 2.**
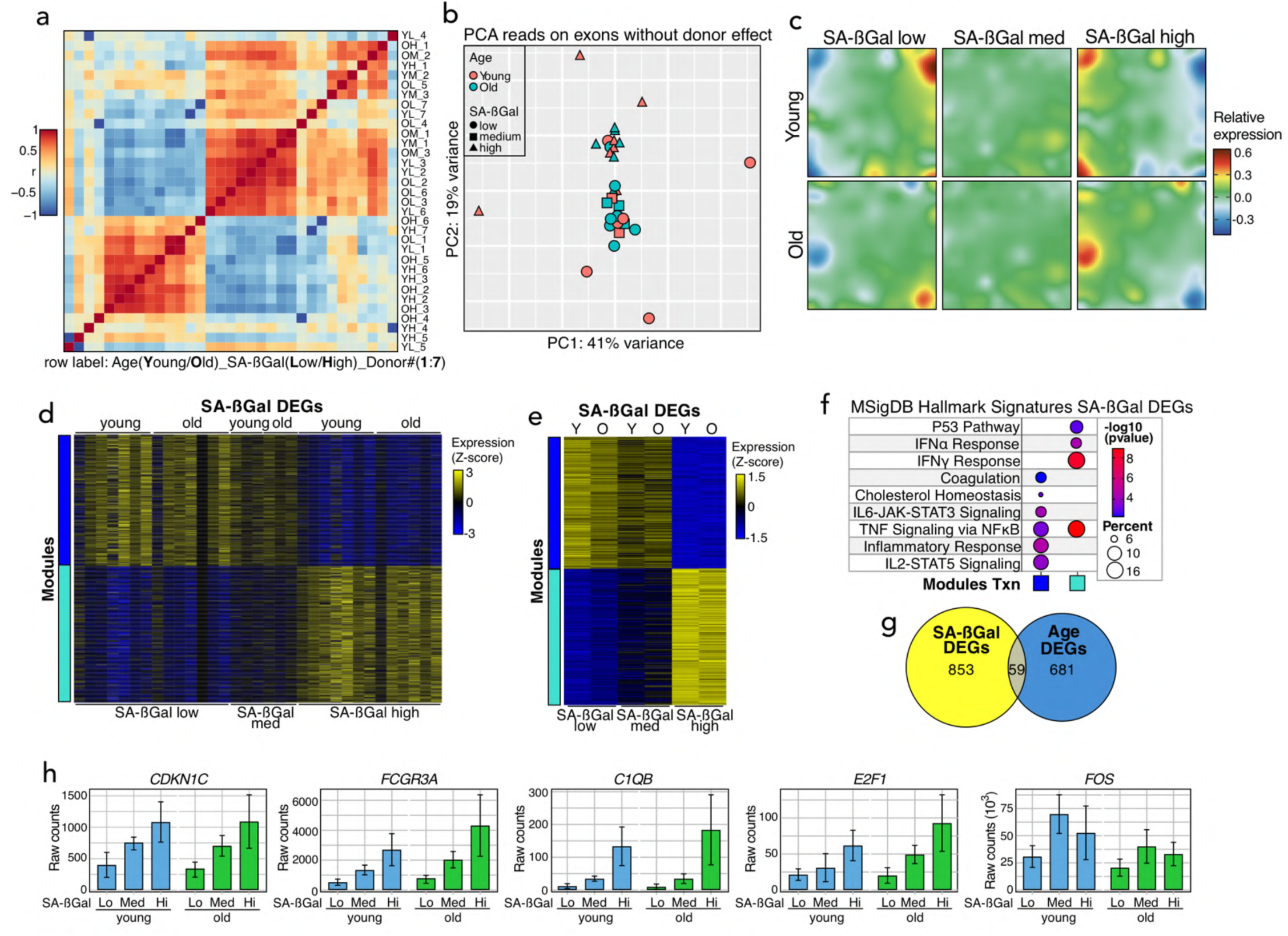
Distinct senescence and aging transcriptional programs in human monocytes. **a.** Correlation heatmap between the transcriptomes of SA-βGal-low, –medium and –high CD14-positive monocytes of older and younger donors after correcting for individual (donor) variation. **b.** PCA projection plot of reads on exons of the indicated transcriptomes of SA-βGal-low, –medium and –high CD14^+^ monocytes from older and younger donors after correcting for donor variation. **c.** Averaged SOMs of donor variation-corrected transcriptomes of SA-βGal-low, –medium and –high CD14-positive monocytes of younger and older donors. **d, e.** Individual (d) and averaged (e) heatmaps of color-coded modules of DEGs in the transcriptomes of CD14-positive monocytes of the indicated SA-βGal activity groups of younger and older donors. The scale bar indicates the row z-score of the rlog-transformed counts. **f.** Functional overrepresentation analysis map showing significant associations of the MSigDB hallmark gene sets for each module described in (d, e). Circle fill is color coded according to the false discovery rate (FDR)-corrected *p*-value from a hypergeometric distribution test. Circle size is proportional to the percentage of genes in each MSigDB gene set. **g.** Euler plot showing the overlaps between SA-βGal– and age-associated DEGs. **h.** Bar plots showing the average raw counts of a subset of genes displaying increased expression in an SA-βGal activity level-dependent manner in the transcriptomes of younger and older donors. Data from the following number of donors: SA-βGal-low, n=7; SA-βGal-med, n=3; SA-βGal-high, n=7 (A-H).

### Conserved transcription factor networks act on a pre-established chromatin landscape to regulate senescence and aging transcriptomes in CD14-positive monocytes

To gain insight into the regulation of senescence and aging gene expression programs in monocytes, we performed the assay for transposase accessible chromatin followed by sequencing (ATAC-Seq)^53^ on sorted CD14-positive monocytes across SA-βGal activity levels from the same 7 younger and 7 older donors used for bulk transcriptome characterization. After controlling for donor and batch effects in an analogous manner to our transcriptome analysis, we did not identify senescence-associated differentially accessible regions (DARs) or a significant correlation to the gene expression output within +/− 500kb from the TSS of senescence DEGs (**Figure 3a**). Given the short half-life of monocytes in blood^16^, we reasoned that these cells, like other innate immune cell types^54,55^, leverage their existing TF networks on a pre-established chromatin landscape to respond quickly to environmental and intrinsic inputs^56^. Consistent with this, TF footprinting and quantification of the TF binding instances of accessible chromatin^+/−^ 500kb from the TSS of senescence-specific DEGs revealed an asymptotic distribution of bound instances dominated by constitutively expressed lineage-determining TFs SPI1, myeloid enhancer factor (MEF) 2A and interferon response factor 1 (IRF1) as well as ZNF384 (**Figure 3b**), a TF often dysregulated in acute leukemia^57^. We then computed TF interactions across SA-βGal activity levels and found widespread, age-independent TF network rearrangements within senescence-associated accessible chromatin (**Figures 3c,d**). To visualize the dynamics and complexity of the TF interactome, we built TF networks using the co-binding interactions and gene expression status of the TFs within the network. As with the senescence TF networks of CD8^+^ T cells^10^, the monocyte senescence TF network was three-layered hierarchical (top, middle, and bottom), composed mostly of constitutively expressed TFs. Surprisingly, and in contrast to CD8^+^ T cells^58^, 70% of the nodes in the monocyte senescence TF network localized to the top layer, with the remaining 30% distributed across the middle (5%) and bottom (25%) layers (**Extended Data Figure 3a**). Given the unusual organization of the network, we evaluated the dynamics, number of bound regions, interactivity (out-degree), and prestimulation of TFs within each layer. This analysis revealed the underlying complexity of the top layer, which displayed specific behaviors for different TF families. Consistent with their role in determining the myeloid lineage, ETS and MEF family TFs were the most pervasively bound and, consequently, mediated most of the interactions with other TFs, irrespective of donor age (**Figures 4a,b**; top layer). Similar behavior was exhibited by AP-1, KLF, and SP family TFs, whereas Homeobox and retinoic acid receptor (RAR) TFs were the most dynamic TFs in this layer. By comparison, the behavior of TFs in the middle and bottom layers, including GATA, REL, and TP53 family members, was more homogeneously dynamic (**Figures 4a,b**; middle and bottom layers). Despite the biochemical heterogeneity of the top-layer nodes, Tarjan’s strongly connected components algorithm^59^ collapsed these TFs into a single, unified regulatory meta-node (**Figures 4c,d**), reflecting the extensive co-binding across shared regulatory regions in the accessible chromatin near senescence-specific DEGs. An analogous integrative analysis of accessible chromatin^+/−^ 500 kb from age-specific DEGs revealed near-identical results. There was no correlation between chromatin accessibility and gene expression output, although we observed a slight, non-significant increase in the chromatin accessibility of monocytes from older donors (**Extended Data Figure 4a;** see also single-cell multiome section). TF footprint quantification and co-binding analysis revealed an age-specific TF network with near-identical properties to that of the senescence TF network: a three-layered network with a highly complex top layer dominated by ETS, MEF and AP-1 TFs which controlled interactions with TFs from comparatively simpler middle and bottom layers (**Extended Data Figures 3b** and **4b-e**). In conclusion, our comprehensive network analysis demonstrates that monocytes regulate leverage their pre-existing TF networks to regulate largely non-overlapping senescence– and age-associated gene expression programs.

**Figure 3.**
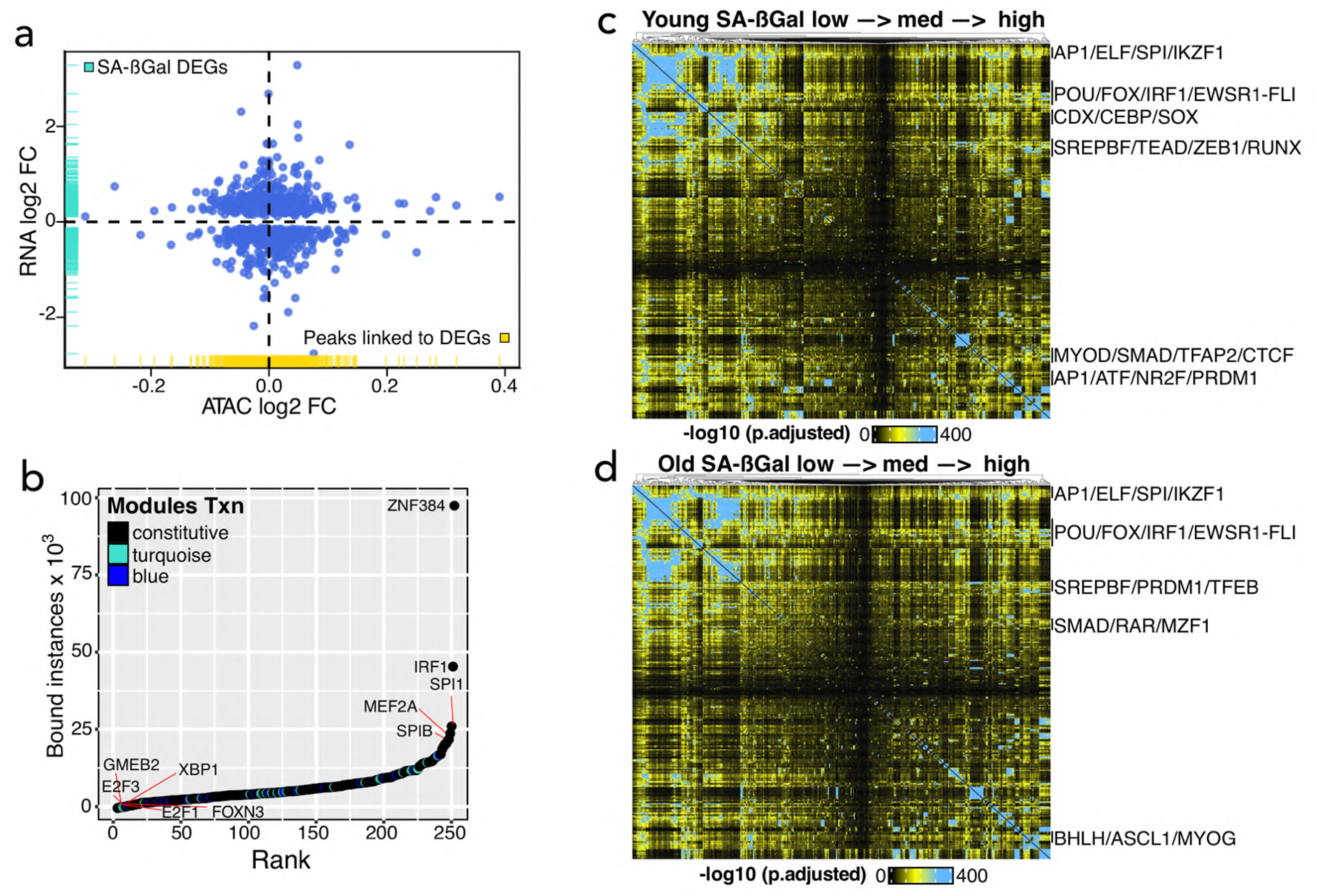
Widespread TF network reorganization during the transitions to senescence and aged monocyte states. **a.** Correlation plot of the RNA log2-fold change of SA-βGal-associated DEGs and the average log2-fold change in the accessibility of their linked peaks (^+/−^ 500kb). Each blue dot represents a DEG linked to at least one chromatin accessibility peak, with its x-axis position reflecting the mean log2-fold change across all peaks linked to that gene. Turquoise ticks on the y-axis represent the RNA log2-fold change distribution of all SA-βGal DEGs. Gold ticks on the x-axis represent the distribution of mean chromatin accessibility log2-fold changes for each SA-βGal DEG. **b.** Rank plot showing the summed binding instances of TFs at SA-βGal DEG-associated chromatin accessibility peaks (^+/−^ 500kb). **c, d.** Co-binding matrices of TF interactions (500 bp resolution; Ward’s criterion for clustering) across SA-βGal activity levels at SA-βGal DEG-associated peaks from younger (d) and older (e) donors. The corresponding q values of the interactions are projected onto the clustering and represented in a color scale defined by their significance using a hypergeometric distribution test. Data from: SA-βGal-low, n=7 donors; SA-βGal-med, n=3 donors; SA-βGal-high, n=7 donors (a-e).

**Figure 4.**
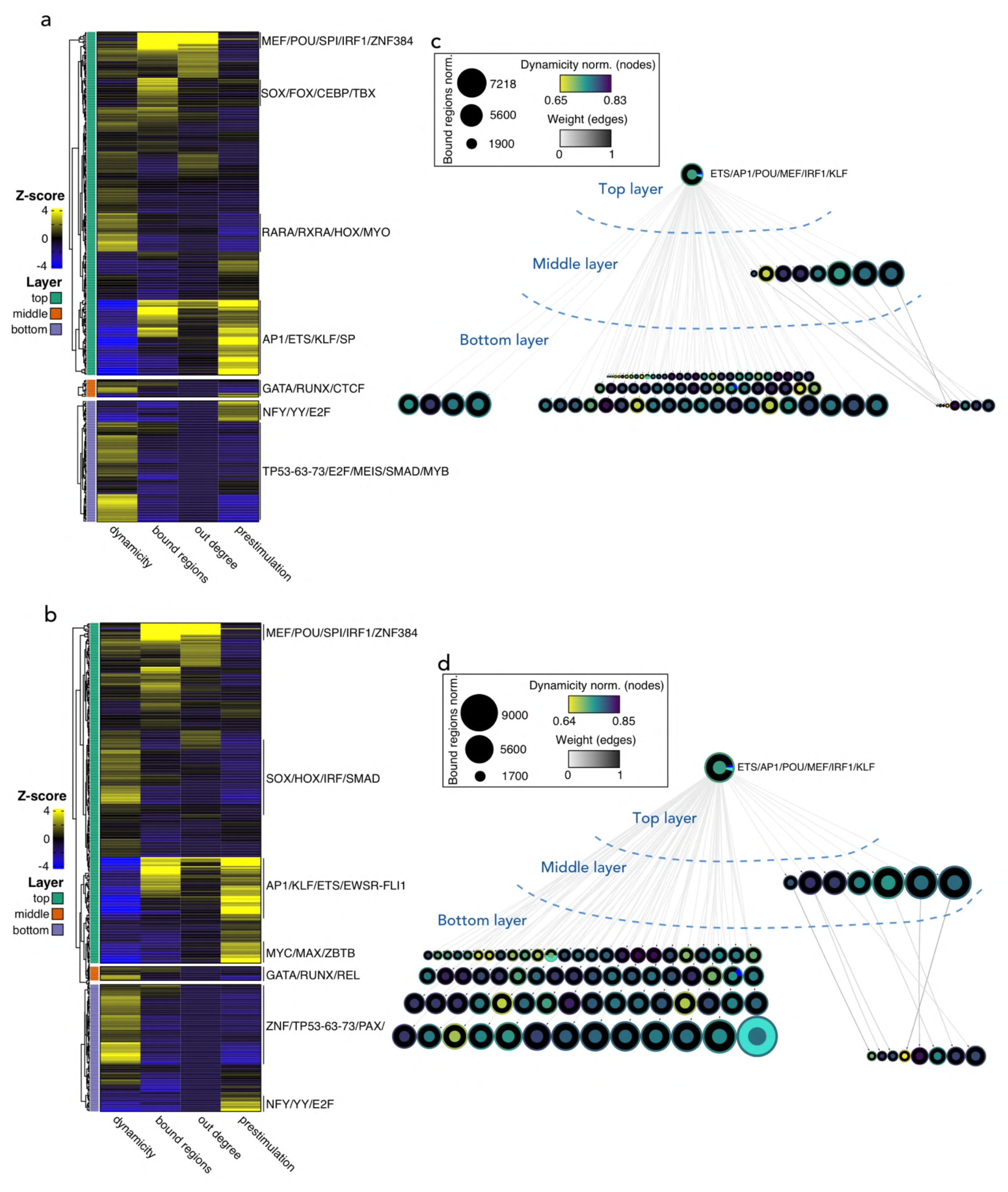
Widespread TF network reorganization during the transitions to senescence and aged monocyte states. **a, b.** Heatmaps showing the variability of TFs in each layer of the networks of younger (a) and older (b) donors across dynamicity, number of bound regions, interactions (out degree) and prestimulation. The scale bar indicates the row Z score of the metric value of each TF motif in the respective network layer. **c, d.** Simplified TF networks at the accessible chromatin (^+/−^ 500kb) of SA-βGal-associated DEGs of younger (c) and older (d) donors. Nodes represent strongly connected components to facilitate visualization. The fill color of the node’s inner circle is based on the normalized dynamicity (prestimulation) of TFs. The fill color of the outer ring indicates whether the TF is constitutively expressed (black) or belongs to a transcriptomic module as described in Figure 1d. The node’s size is proportional to the bound regions by a given TF(s). Each network has three layers: (1) the top layer with no incoming edges, (2) the core layer with incoming and outgoing edges, and (3) the bottom layer with no outgoing edges. The inset denotes the bound regions scale, prestimulation and directional overlap of the TF interactions. Networks were constructed from chromatin accessibility data from: SA-βGal-low, n=7 donors; SA-βGal-med, n=3 donors; SA-βGal-high, n=7 donors.

### A transitional senescence program leads to an age-associated inflammatory monocyte state

Our flow cytometric and bulk multiomic profiling analyses suggested the existence of two CD14-positive monocyte populations defined by senescence– and age-associated gene expression patterns. To define the single-cell heterogeneity and dynamics of senescence– and age-associated monocytes during human aging, we performed single-cell (sc) multiomic profiling (single-nucleus transcriptome and chromatin accessibility) on magnetic bead-enriched CD14-positive monocytes from 4 young and 3 older donors using the 10X Genomics platform, obtaining 8,758 high-quality cells. Integration of the scRNA-seq datasets resulted in homogenous cell mixing across samples and a UMAP architecture consisting of two main monocyte populations connected by a ‘bridge’ (**Figures 5a,b**). Clustering analysis identified 6 cell clusters, 0 to 5, which revealed two main monocytes states: a resting, non-inflammatory state (clusters 2 and 4) and an inflammatory state (clusters 0, 1, 3 and 5) that enriched for senescence and aged monocyte gene signatures derived from our bulk transcriptome analysis (**Figures 5c,d** and **Extended Data Figures 5a,b**). Quantification of cluster proportions revealed an expansion of inflammatory clusters 0, 1 and 5 with a concomitant reduction of non-inflammatory clusters 2 and 4 in monocytes from older individuals (**Figure 5d**). To confirm the identity of the cell populations in our dataset, we performed label transfer using the Azimuth PBMC reference^60^. This approach identified ∼80% of cells as CD14^++^CD16^-^ classical monocytes, which comprised most of the two major monocyte states in the UMAP. Notably, the ‘bridge’ connecting these two populations was a mixture of CD14^++^CD16^-^ classical and CD14^+^CD16^++^ non-classical monocytes (∼12.3% of all cells), indicating a transitional population of double-positive monocytes (**Extended Data Figure 5c**). To identify ‘young’ and ‘old’ monocyte markers in our dataset, we determined age-specific DEGs. We identified 798 young– and 228 old-specific DEGs that defined basal/resting and inflammatory monocyte states, respectively (**Figures 5e**,**f**). Visualization of the top 100 young and old DEGs at single-cell resolution revealed a shift in the magnitude of the gene expression output, suggesting that fine-tuning of the transcriptional output, rather than a binary switch, underlies this monocyte state transition (**Figure 5g**). To locate young and aged monocytes, we calculated and subsequently projected their DEG module score (scYoung and scOld, respectively) onto the UMAP. Consistent with a cell state transition, the scYoung module enriched almost exclusively on non-inflammatory monocytes in clusters 2 and 4 whereas the scOld module displayed a gradient of increasing expression from cluster 0 through the bridge and achieving its maximum in monocytes in clusters 1 and 3 (**Figures 5h,i**). To validate these results, we projected our age (AgeSig) and senescence (SenSig) bulk gene signatures. Expectedly, the expression of AgeSig displayed a similar distribution to the scOld module whereas, strikingly, SenSig was maximally expressed by the double-positive cluster 5 monocyte population that comprised the bridge (**Figures 5j,k**). Importantly, we verified an overrepresentation of bulk senescence and aged monocyte signatures in older donors (**Extended Data Figures 5d,e**). To provide further evidence of a transitional senescence state, we performed pseudotime analysis using Monocle3^61^. Using the ‘youngest’ monocyte as origin, we observed a natural trajectory towards the aged monocyte state via the senescence bridge, which we corroborated with an unbiased RNA velocity analysis^62^ (**Figures 6a,b**). Furthermore, the transcriptional entropy increased in a pseudotime-dependent manner, reaching near-maximum levels at the senescence bridge, before stabilizing at the inflammatory monocyte state (**Figures 6c-e**). This increase in transcriptional entropy is reminiscent of fibroblasts undergoing oncogene-induced senescence (OIS)^42^. Collectively, the results from our comprehensive single-cell transcriptome analysis provide strong evidence for a transitional senescence monocyte state that expands during natural human aging.

**Figure 5.**
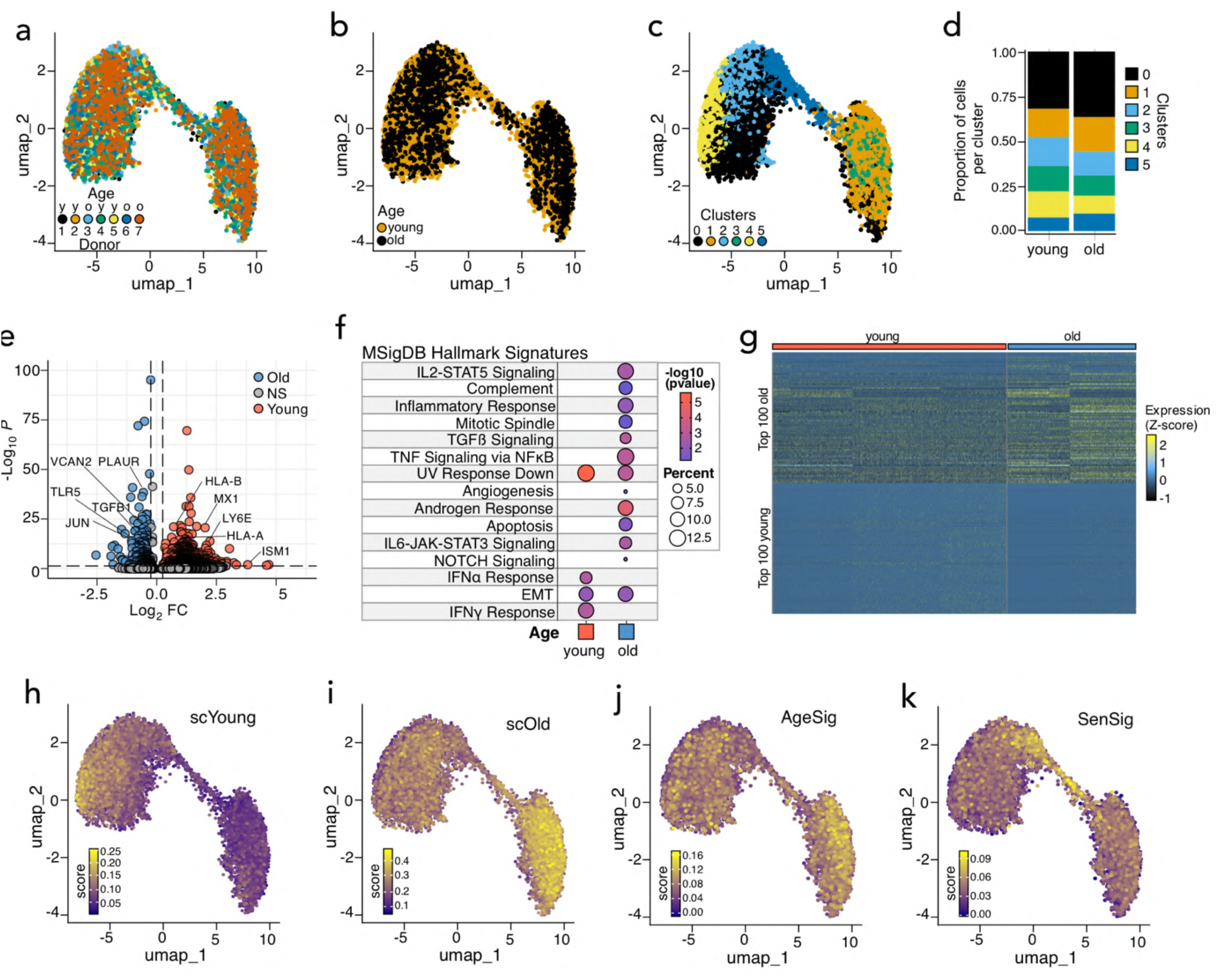
A transitional senescence state leads to an age-associated inflammatory monocyte state. **a, b.** UMAPs of the integrated single-cell gene expression data (8,758 monocytes) of the indicated donors (a) and age groups (b). **c, d.** UMAP showing the location of the identified Seurat clusters (c) and their redistribution across age groups. For d, all age-associated cluster shifts were significant at *p* < 6.0995 x 10^-4^, Fisher’s exact test. **e.** Volcano plot showing the age-associated DEGs in the dataset. Age-specific DEGs are color-coded. **f.** Functional overrepresentation analysis map showing significant associations of the MSigDB hallmark gene sets for each age-associated DEG set from e. Circle fill is color coded according to the false discovery rate (FDR)-corrected *p*-value from a hypergeometric distribution test. Circle size is proportional to the percentage of genes in each MSigDB gene set. **g.** Heatmap showing the single-cell expression of the top 100 age-associated DEGs from e. **h-k.** UMAPs showing the UCell enrichment score and location of young-(h) and old-(i) specific DEGs from e as well as the age (AgeSig) and senescence (SenSig) gene signatures derived from bulk RNA-seq sets from Figure 2 and Extended Data Figure 2.

**Figure 6.**
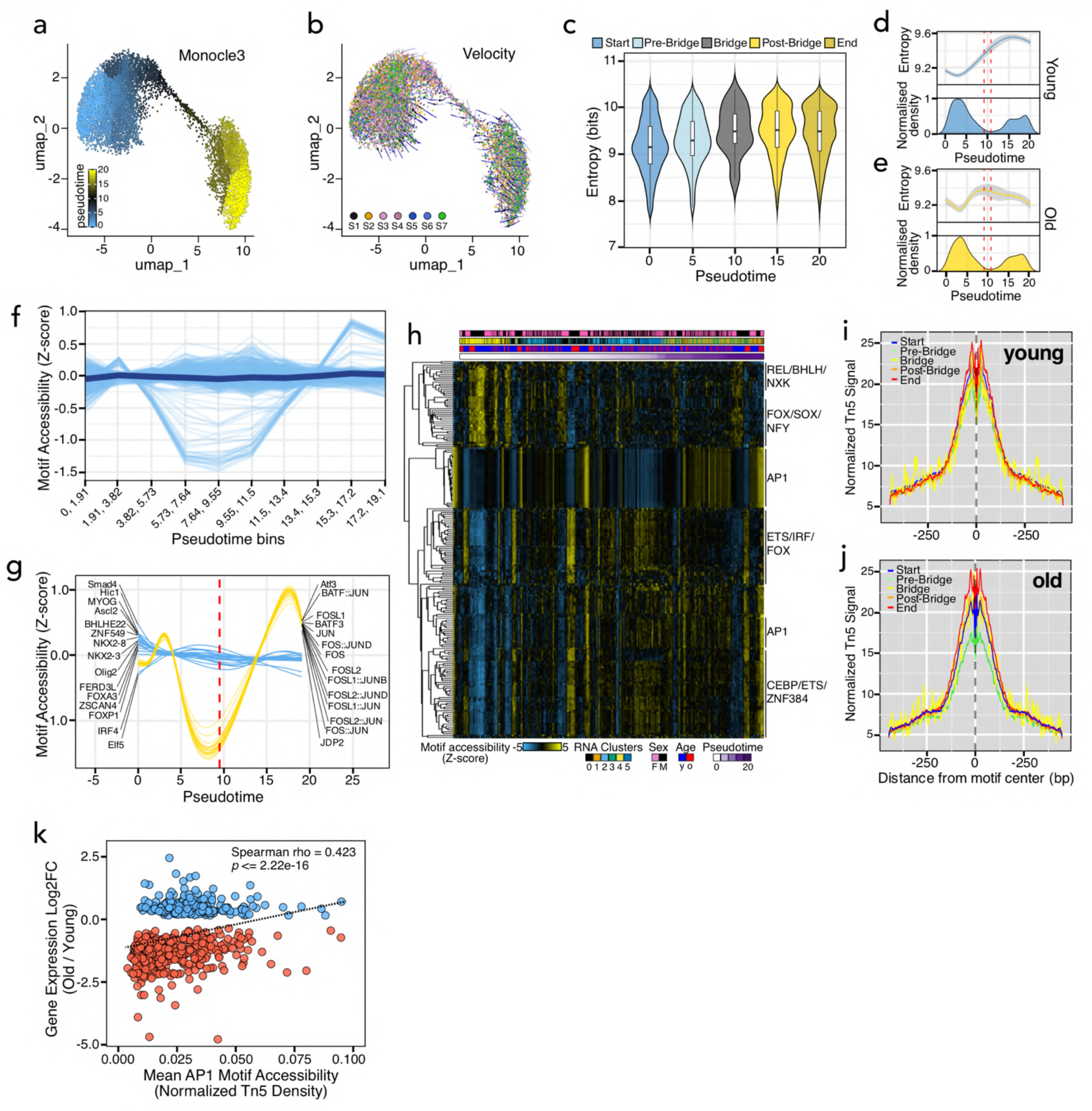
A transitional senescence state leads to an age-associated inflammatory monocyte state. **a, b**. UMAPs of the integrated single-cell gene expression data showing Monocle3 pseudotime (a) and RNA velocity (b) trajectories. For a, the cell with the most negative UMAP-1 dimension value was chosen as root cell. **c.** Violin plots showing the distribution of the transcriptional Shannon entropy across pseudotime bins. **d, e.** Combined entropy and population density dynamics plot of monocytes from young (e) and old (e) donors. The entropy lines were smoothed using a generalized additive model (GAM) smooth regression with 95% confidence intervals. The vertical dashed lines indicate the transition bridge. **f.** Raw chromVAR motif accessibility dynamics for the top 200 variable TFs along pseudotime. Thin light blue lines represent individual TF motif accessibility scores while the thick dark blue represents the average of all 200 motifs. **g.** As in f but showing the smoothed motif accessibility scores for the top 20 active TFs in monocytes from young and old donors. Trajectories were smoothed using a GAM smooth regression. **h.** Heatmap showing the motif accessibility dynamics of the top 200 variable TFs along pseudotime at single-cell resolution. The top annotations are defined at the bottom of the heatmap. A subset of TF families with dynamic behavior before and after the bridge is indicated to the right of the heatmap. A rolling average of *k*=50 was used to smooth the signal in the heatmap. **i, j.** Footprinting metaprofiles of AP-1 family dimers (FOS::JUN, FOSL2::JUN, FOSL2::JUNB, FOS::JUND, FOS::FOSL2, FOSL2::JUND, FOS::JUNB, FOSL1::JUNB, FOSL1::JUND, JUN, FOSL2 and FOSL1) along pseudotime at accessible chromatin^+/−^500kb of CD14^+^ monocyte age-specific DEGs (single-cell). **k.** Scatter plot showing the correlation between increased local AP-1 motif accessibility and expression of age-associated DEGs (cut-off of absolute 0.25 log2 fold-change in at least 10% of cells). The rho and *p*-value of a Spearman correlation analysis are shown.

### AP-1 regulatory dynamics and global chromatin remodeling orchestrate senescence and aging monocyte transitions

To dissect the gene-regulatory mechanisms underlying this monocyte state transition, we sought to identify age-related changes to the accessible chromatin at single-cell resolution. Consistent with the results from our bulk analysis, there was no correlation between chromatin accessibility and gene expression dynamics at the single-cell level. Similarly, we observed a slight, non-significant increase in the chromatin accessibility of monocytes from older donors (**Extended Data Figure 6a**). A granular fragment length distribution analysis of a 10 megabase-(mb) long region of chromosome 1 revealed that the increased accessibility of chromatin of monocytes from older donors was characterized by an unusually high number of sub-nucleosomal fragments (**Extended Data Figure 6b**). This fragment size distribution was consistent genome-wide and across pseudotime at both promoters and distal regions (**Extended Data Figures 6c-e**). These results strongly suggest a global rearrangement of chromatin architecture during monocyte aging. In parallel, we profiled the TF network to identify age-specific activity signatures. Quantification of TF footprints of pseudobulked chromatin accessibility profiles of individual donors revealed a binding distribution consistent with that observed in bulk data (**Extended Data Figure 6f**). Clustering of pseudobulked differential TF binding activity profiles revealed age-specific patterns, as monocytes from older donors exhibited high activity of AP-1, IRF and SPI family TFs whereas monocytes from younger donors displayed preferential use of FOX and SOX TFs, although there were some donor-specific differences (**Extended Data Figure 6g**). These age-specific TF activity profiles were largely reproducible at single-cell resolution using an alternative method that relies on motif accessibility rather than direct footprinting^63^ (**Extended Data Figure 6h**). To identify the key TFs driving the transition towards the inflammatory monocyte state, we plotted the motif accessibility of the top 200 most variable TFs along pseudotime as a proxy for their binding. While the activity of most of these TFs remained constant along the pseudotime trajectory, a distinct subset, most notably AP-1 family TFs, exhibited transitional kinetics. Specifically, their activity dropped sharply upon approaching the transitional senescence bridge followed by a rapid surge as the monocytes transitioned towards the inflammatory state (**Figures 6f,g**). This regulatory transition, characterized by increased activity of additional pro-inflammatory TF families such as ETS, IRF and C/EBP, occurred independently of donor sex and age (**Figure 6h**). To confirm the inferred AP-1 binding dynamics, we analyzed single-cell TF footprint metaprofiles across AP-1 motifs within +/− 500kb of age-specific DEGs along pseudotime. We observed progressive AP-1 binding along the trajectory, reaching peak occupancy at the inflammatory monocyte state in both young and older donors (**Figures 6i,j**) which was positively correlated with increased expression of age-specific DEGs (**Figure 6k)**. This increased AP-1 binding activity was confirmed by quantifying and comparing the footprints of JUN as well as of lineage-determining factors SPI1 and IRF1 along pseudotime and across all accessible chromatin (**Extended Data Figures 6i-k**). In summary, our comprehensive single-cell multiomic analysis identifies AP-1 TFs as critical mediators of senescence and inflammatory monocyte state transitions, highlighting AP-1-dependent signaling as a potential target for therapeutic intervention in systemic aging.

### Accumulation of inflammatory, pre-septic monocytes in aging humans

Aging increases the basal expression of inflammatory mediators in both the innate and adaptive compartments, a process known as inflammaging. This higher inflammatory baseline is associated with increased susceptibility to and impaired resolution of infection, which in turn results in an increased risk of adverse outcomes such as sepsis^6,64,65^. To investigate the potential contribution of the senescent and aged monocyte populations to clinical complications, we integrated our scRNA-seq dataset with a comprehensive reference of monocyte transcriptomes across various sepsis cohorts of distinct severities^66^. UMAP visualization of the integrated datasets revealed a preferential co-localization of young monocytes with control and low-severity septic (Leuk-UTI) monocytes whereas old monocytes were mostly localized with monocytes of more severe disease states including intermediate urosepsis (Int-URO; URO), non-septic and septic intensive care unit admission (ICU-NoSEP; ICU-SEP) and bacterial sepsis (Bac-SEP) (**Figure 7a** and **Extended Data Figure 7a**). Projection and quantification of the expression of our single-cell old (scOld) and senescence (c5) signatures revealed a progressive increase in the number of cells positive for both signatures, which correlated with the severity of the sepsis cohort (**Figures 7b-d**). In this context, four distinct monocyte states (inflammatory MS1-3 and non-inflammatory MS4) were recently described that associate with sepsis severity. Specifically, an expansion of a highly inflammatory MS1 state was found in sepsis and sterile ICU cohorts whereas the non-inflammatory MS4 state was the most abundant monocyte state in control individuals^66^. We therefore reasoned that monocytes from older individuals could exhibit an overrepresentation of sepsis-associated transcriptomes. After confirming the spatial distribution, gene expression profile and enrichment of senescence and age signatures across monocyte states from sepsis cohorts (**Extended Data Figures 7b-e**), we quantified the proportion of young and monocytes using a k-nearest neighbor (*k*-NN) classification approach. We observed a ∼2.2-fold increase of monocytes from older donors within the MS1 state, accompanied by a relative decrease in the remaining states compared to younger donors (**Figures 7e-g)**, which was confirmed by calculating the Euclidean distance to the MS centroids (**Figure 7h**). Quantification of the scOld and c5 signatures across monocyte states revealed a progressive increase from the non-inflammatory MS4 to the MS1 state and highlight the MS3 state as the potential senescent transitional state in the sepsis setting, respectively (**Extended Data Figures 7f,g**). Furthermore, we consistently found overrepresentation of our senescence and age transcriptomic signatures in the inflammatory, sepsis-associated gene expression modules in a cohort of patients undergoing or having recovered from sepsis^67^ (**Extended data Figures 7h-j**) as well as in a separate cohort of monocytes from septic, critically ill and cancer patients^68^ (**Figure 8a-g**). Taken together, these results demonstrate that monocytes from older individuals are transcriptionally primed towards sepsis-associated states and highlights a potentially critical role for a transitional senescent monocyte state in driving susceptibility to sepsis.

**Figure 7.**
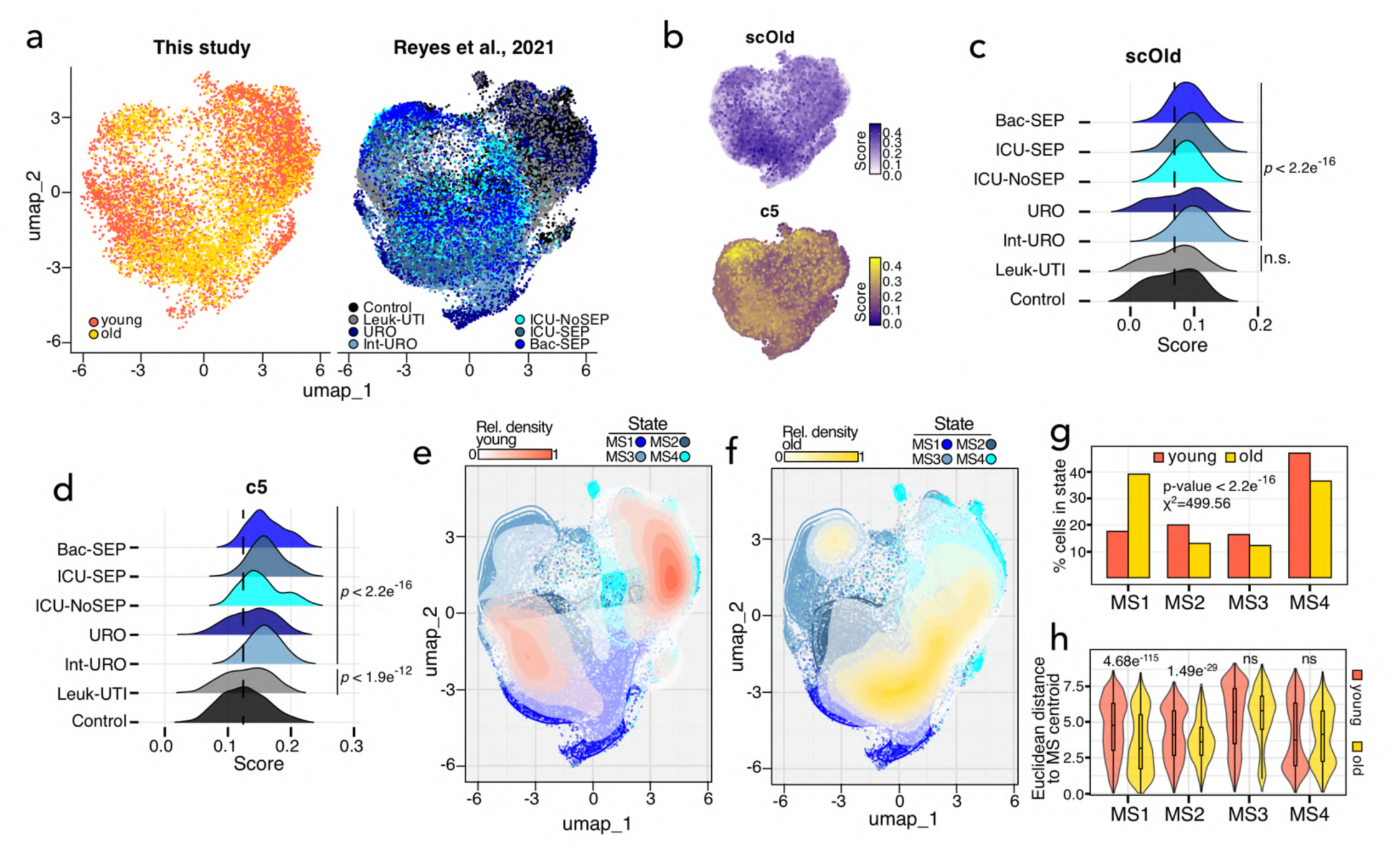
Accumulation of septic monocytes in aging humans. **a.** Integrated UMAP showing the spatial distribution of young and old monocytes (this study) relative to control and septic monocytes from the indicated sepsis cohorts from reference^66^. **B.** Projection of aged (scOld) and senescence (c5) gene signatures identified from the single-cell RNA-seq analysis described in Figures 5c-j and Extended Data Figures 5a, b onto the integrated UMAPs from 7a. Scale bar denotes the UCell score. **C**, **D.** Ridge plots showing the shift of the scOld and c5 gene signature scores across control and septic monocytes from reference^66^. The dashed black lines indicate the median score of each gene signature in control monocytes. Statistical significance of gene signature score shifts in sepsis cohorts relative to control was calculated using post-hoc Dunn tests. **e**, **f.** Contour plots showing the spatial alignment of young (e) and old (f) monocytes within monocyte states defined in^66^. **g.** Histogram showing the quantification of enrichment of young and old monocytes within monocytes states as in e, f. Quantification was performed using k-nearest neighbor (n=10) using young and monocytes as queries and monocyte states as references. Statistical significance was calculated with a chi-squared test, and the *p*-value is shown. **h.** Violin plots of spatial similarity analysis showing the Euclidean distance of young and old monocytes to the centroid of monocyte states in the UMAPs. Statistical significance of the indicated comparisons was calculated using a Wilcoxon rank test.

**Figure 8.**
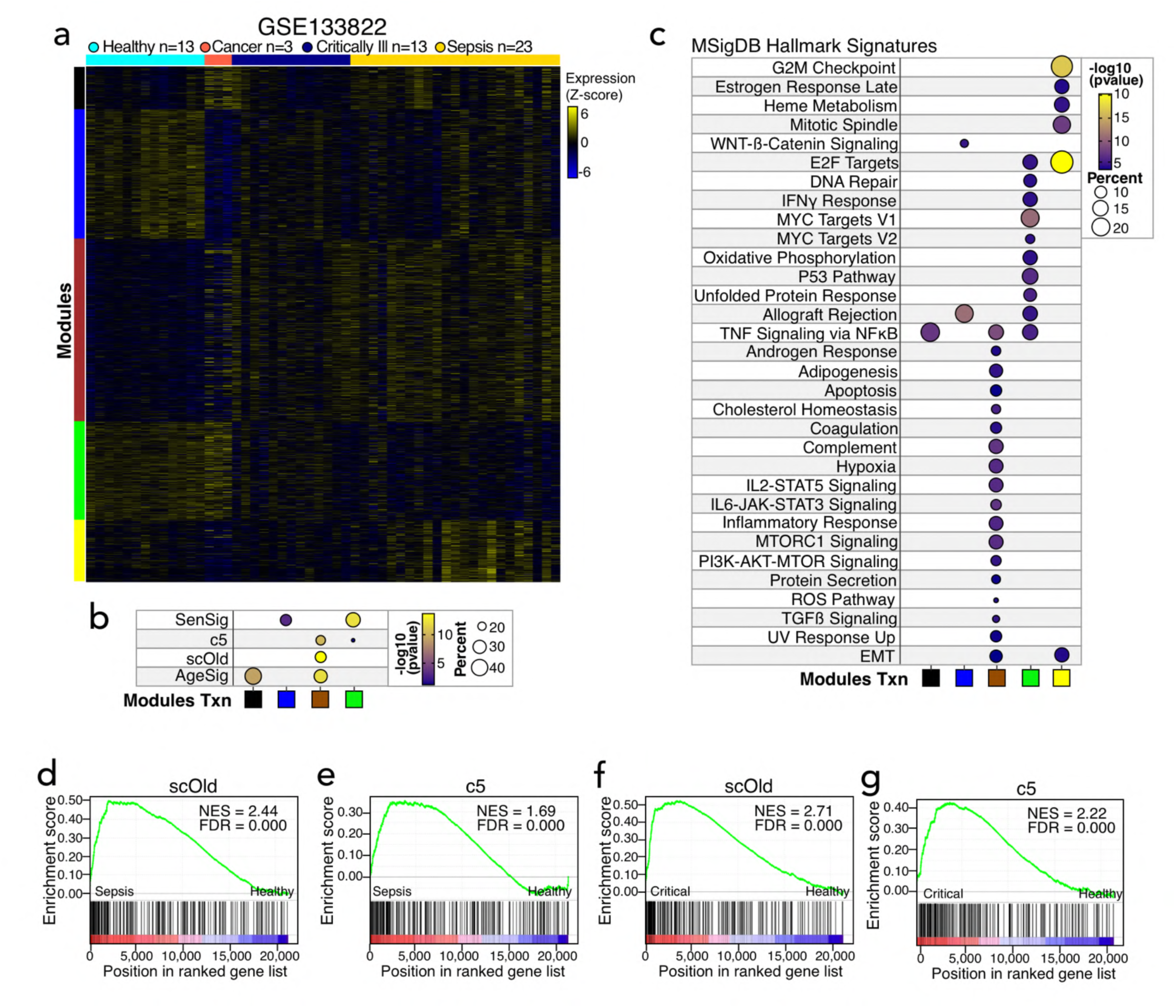
Senescence and age monocyte transcriptional signatures predict sepsis and critical illness. **a.** Individual heatmaps of color-coded modules of DEGs in the transcriptomes of monocytes of patients in the indicated conditions from dataset GSE133822. The scale bar indicates the row z-score of the rlog-transformed counts. **b, c**. Functional overrepresentation analysis map showing significant associations of the indicated age and senescence signatures (b) and the MSigDB hallmark gene sets (c) for each DEG module from h. Circle fill is color coded according to the false discovery rate (FDR)-corrected *p*-value from a hypergeometric distribution test. Circle size is proportional to the percentage of genes in each MSigDB gene set. **d-f.** Gene set enrichment analyses (GSEAs) showing normalized enrichment score (NES) plots and false discovery rate (FDR) values for the aged (scOld) and senescence (c5) gene signatures identified from the single-cell RNA-seq data of Figure 5 in the transcriptomes of septic (d, e) and critically ill (trauma; f, g) patients relative to healthy controls.

## Discussion

The accumulation of senescent cells in tissues is a major driver of age-related pathologies, mediated in large part by local inflammation and tissue microenvironment alterations induced by the SASP^3^. Paradoxically, the immune system-mediated clearance of senescent cells, a process facilitated by the SASP along with expression of major histocompatibility complex (MHC) and NKG2D ligands on tissue-resident senescent cells, becomes progressively impaired with age^5^. Recent evidence points to an aged immune system as an underlying driver of the age-associated accumulation of senescent cells in tissues. While concerted efforts have focused on defining dysfunctional cell states in the adaptive immune compartment during human aging, with a specific focus on CD4^+^ and CD8^+^ T cells^11,69,70^, our understanding of the age-associated dysregulation of the components of the innate immune system is only emerging^65^. Specifically, within the monocytic lineage, recent work has characterized age-associated defective TLR-mediated transcriptional responses^36^ and elevated basal secretion of inflammatory mediators^33^ that are associated with persistent local inflammation^71^. Our current understanding of monocyte populations supports the presence of three subtypes, classical CD14^++^CD16^-^, intermediate CD14^++^CD16^+^ and non-classical CD14^+^CD16^++^, each with their own phenotypic profile and function^72^. However, recent evidence shows that, within these three subtypes, there is increased transcriptional and functional heterogeneity in homeostasis as well as in disease states^66,67,73^. Importantly, the underlying gene-regulatory mechanisms that drive inflammation– and disease-associated monocyte state transitions are poorly understood. Elucidating the epigenetic mechanisms driving disease-associated monocyte states holds potential for developing targeted therapies to restore monocyte function, potentially mitigating systemic inflammation while enhancing responses to infection.

Given the outsized role of monocytes as early sensors of infection and tissue damage as well as orchestrators of adaptive immunity^17^, and the recently documented effect of ageing on circulating monocyte populations^74^, we utilized a multidimensional, high-resolution integrative approach to define age– and senescence-associated monocyte transitions. Our findings uncover a transitional monocyte state with senescence features, including increased SA-βGal activity and upregulation of CDKN1A/p21, CDKN1B/p27 and JUN, that fuels the accumulation of inflammatory monocytes in older humans, establishing the core gene-regulatory principles underlying monocyte-driven inflammaging and revealing potential inroads for targeting systemic inflammation. Our data show that this heterogeneous transition, originating within the major CD14^++^CD16^-^ classical population, initiates senescence-associated reprogramming characterized by the emergence of a transitional double-positive population with increased transcriptional entropy and reorganized chromatin architecture. Rather than representing a terminal state, RNA velocity and trajectory analyses demonstrate that this age-associated senescent state is canalized towards an epigenetically distinct, inflammatory monocyte population. Importantly, integration with transcriptomic data from clinical sepsis cohorts^66^ shows that both the age-associated transitional senescent and end-point inflammatory monocytes are primed towards sepsis-associated states. Intriguingly, transient expansions of the CD14^++^CD16^+^ double-positive monocyte subset have also been documented in Crohn’s disease^75,76^, Graves’ disease^77^ and rheumatoid arthritis^78^. Therefore, our results not only highlight the underappreciated epigenetic heterogeneity of circulating monocytes but also emphasize a potentially instructive role of transitional senescence states in determining disease-associated monocyte fate.

We show that master senescence regulators of the AP-1 TF family^42^ play a central role in orchestrating the transition to the inflammatory monocyte state. Mechanistically, AP-1 TFs, in conjunction with myeloid-lineage determining TFs from the SPI and MEF families, leverage a pre-established chromatin landscape to rewire a pre-bound, highly complex network of predominantly constitutive TFs to activate senescence– and age-associated inflammatory gene expression. This unusual network architecture, which likely represents a conserved mechanism to maximize rapid responses to incoming stimuli in short-lived cells^54,55^, contrasts with our previous work on long-lived fibroblasts and CD8^+^ T cells that exhibit a higher dependency on active chromatin remodeling and inducible expression of TF network nodes^42,43,58^. Interestingly, chronic TLR4 stimulation of bone marrow-derived monocytes leads to an expansion of a sepsis-associated monocyte state through a transcriptional program that is dependent on a gene-regulatory network comprising AP-1, C/EBP and SPI1 TFs^66^. Given the potentially outsized impact of disease-associated monocyte states on clinical outcome^40^, disruption of AP-1-mediated transcriptional programs emerges as a highly attractive therapeutic strategy to modulate monocyte senescence and systemic inflammation.

In conclusion, the present work establishes the gene-regulatory principles governing monocyte senescence-associated transitions and offers a logical framework to modulate systemic inflammation. It also emphasizes the potential of multidimensional integrative profiling as a powerful approach to uncover previously unknown senescence-associated cell fate transitions.

## Methods

### Study participants’ details

The Institutional Review Board of New Jersey Medical School approved this study (ID Pro2020000006, approved May 14, 2024). Peripheral blood mononuclear cells (PBMCs) were obtained from the blood of consenting healthy donors collected in heparinized tubes (BD Biosciences, Franklin Lakes, NJ) and used for flow cytometric analyses. Leukopacks were commercially obtained from the New York Blood Center for bulk and single-cell high-throughput sequencing studies.

### Peripheral blood mononuclear cell (PBMC) isolation and CD14-positive monocyte enrichment

PBMCs were isolated using density gradient centrifugation with lymphocyte separation medium (Corning; 25-072-CV). Heparinized blood was diluted 1:1 with Hanks’ Balanced Salt Solution (HBSS), (Corning; 21-020-CM), layered over lymphocyte separation medium at a 2:1 ratio, and centrifuged at 400 x *g* for 30 min with reduced deceleration (deceleration factor of 2). CD14-positive monocytes were enriched from PBMCs buffy coat using positive selection using magnetic human CD14 Microbeads and LS columns (Miltenyi; 130-050-201 and 130-042-401), following the manufacturer’s protocol. The purity of the isolated CD14^+^ monocytes was assessed via flow cytometry by staining for CD14 (APC-conjugated anti-CD14 REA monoclonal antibody; Miltenyi; 130-110-578) 4°C for 20 min in 1x phosphate-buffered saline (PBS) pH 7.2 ^+^ 2% fetal bovine serum (FBS; Gibco, 26140-079) while protected from light. Samples were subsequently washed 1x PBS^+^ 2% FBS and kept on ice before acquisition on a BD LSRFortessa X-20 (BD Biosciences). Purity of samples was ∼90% across isolations.

### SA-βGal staining, FACS and cyCONDOR analysis

PBMCs or enriched CD14-positive monocytes were resuspended in RPMI-1640 medium containing 10% FBS and 1× penicillin/streptomycin (Corning; 30-002-Cl). The Cellular Senescence Detection Kit-SPiDER-βGal (Dojindo Molecular Technologies; SG04-10) was used to identify senescent CD14-positive monocytes as we previously described^11^. Briefly, PBMCs or CD14-positive monocytes were incubated with a 1:1000 dilution of bafilomycin A-1 for 30 min before the addition of 1:1000 dilution SPiDER-βGal for an additional 30 min all whilst at 37°C in a 5% CO_2_ incubator. SA-βGal stained cells were subsequently washed with 1x PBS and resuspended in 1x PBS ^+^ 2% FBS for surface staining using an APC-conjugated anti-CD14 antibody as described above. Cell sorting on CD14-positive monocyte-enriched preparations was performed on a BD FACSAria Fusion instrument based on 4′,6-diamidino-2-phenylindole (DAPI)-negative (BioLegend; 422801) cells, collecting SA-βGal-low (bottom 30%), SA-βGal-med (middle 30%) and SA-βGal-high CD14-positive (top 30%) monocytes using older donors as a reference. For conventional flow cytometry, 1×10^6^ PBMCs were stained with BUV395-conjugated anti-CD45 (BD Biosciences, 563792), APC-conjugated anti-CD3 (BioLegend, 300312), Brilliant Violet 421-conjugated anti-CD56 (BioLegend, 632552), Alexa Fluor 700-conjugated anti-CD19 (BioLegend, 302226) and APC/Cyanine7-conjugated anti-CD14 (BioLegend, 367108). For cyCONDOR analysis, 1×10^6^ PBMCs were stained with BV605-conjugated anti-CD16 (3G8), Pac Blue-conjugated anti-CD19 (HIB19), PerCP-conjugated Cy5.5 anti-CD3 (HIT3a), PE-Cy7-conjugated anti-HLA-DR (Tü36), APC-Cy7-conjugated anti-CD14 (63D3) (BioLegend). Simultaneously, PBMCs were stained with MT1 MitoMP, MitoBright LT Deep Red and Lipi-Red (Dojindo Molecular Technologies; MT13-10, MT12-12 and LD03-10) at 8000X, 10000X and 1500X dilutions respectively. Fluorescence acquisition was performed on a BD Symphony instrument. Two separate acquisitions of 12 (6 young and 6 old; MT1 MitoMP and MitoBright LT Deep Red) and 16 (8 young and 8 old; Lipi-Red) were analyzed using cyCONDOR after export from FlowJo as recommended by the developers. 5,000 random cells from each donor were extracted, auto-logically transformed and batch-corrected (day of experiment) using Harmony. Quantification of monocyte subsets was performed using a custom script in R studio using base R functions, dplyr and ggplot2. UMAP projections and scaled marker expression plots were generated using the standard cyCONDOR workflow and statistical significance for marker expression distributions was performed using the Wilcoxon test.

### RNA-seq

RNA from SA-βGal-low (n=7 per age group), SA-βGal-med (n=3 per age group) and SA-βGal-high CD14-positive (n=7 for young, n=6 for old) monocytes was purified using an RNA XS Plus kit (Macherey-Nagel; 740990.250) according to the manufacturer’s instructions). RNA integrity was evaluated using a TapeStation 4200 system (Agilent), and only RNA with an integrity of number of ≥ 7 was used for library preparation. Libraries were constructed using the NEBNext Ultra II Directional RNA Library Prep Kit (with Poly(A) mRNA isolation) (New England Biolabs; E7760L) according to the manufacturer’s instructions. Paired-end sequencing was performed on an Illumina NovaSeq X instrument. At least 40 million reads per sample (20 million per strand) were obtained and used for downstream analyses.

### ATAC-seq

The transposition reaction and library construction were performed as described in^53^. Briefly, 50,000 SA-βGal-low, SA-βGal-med and SA-βGal-high CD14-positive monocytes were washed in 1× in PBS, and centrifuged at 500 × *g* at 4°C for 5 min. Nuclei were extracted by incubating cells in nuclear extraction buffer (containing 10 mM Tris-HCl, pH 7.4, 10 mM NaCl, 3 mM MgCl2, 0.1% IGEPAL CA-630) and immediately centrifuged at 500 × *g* at 4°C for 5 min. The supernatant was removed by pipetting, and the transposition was performed by resuspending nuclei in 50 μl of Transposition Mix containing 1× TD Buffer and 2.5 μl Tn5 transposase (Illumina; 20034211) for 30 min at 37°C. DNA was extracted using a MinElute kit (Qiagen; 28006). Libraries were produced by PCR amplification (8 cycles) of tagmented DNA using an NEB Next High-Fidelity 2× PCR Master Mix (New England Biolabs; M0544S). Library quality was assessed using a TapeStation 4200 system (Agilent). Paired-end sequencing was performed on an Illumina NovaSeq X instrument. Typically, 30-50 million reads per sample were obtained and used for downstream analyses.

### Single-cell multiome CD14-positive nuclei preparation

CD14-positive monocytes from 4 young donors and 3 old donors were isolated via positive selection from freshly prepared PBMCs using magnetic human CD14 Microbeads and MACS LS columns in a QuadroMACS magnetic separator (Miltenyi; 130-050-201, 130-042-401 and 130-042-401) according to manufacturer’s instructions. Briefly, ∼100 million PBMCs per donor were resuspended in 400 μl buffer (1X PBS-pH 7.2, 0.5% FBS and 2mM EDTA) and incubated with 100 μl CD14 microbeads for 15 minutes at 4°C. PBMCs were then washed with 10 ml buffer, pelleted at 300 x *g* for 10 minutes and resuspended in 500 μl buffer. PBMC suspension was applied onto LS columns placed on the magnetic separator, flow-through containing unlabeled cells was discarded, columns were washed 3 times with buffer and columns were removed from magnet and CD14-positive cells were immediately eluted in a collection tube using 5 ml of buffer. CD14^+^ monocytes were manually counted using a hemocytometer and viability was assessed via 0.4% Trypan blue staining (>= 85%). One million freshly isolated monocytes per donor were processed the same day for generation of single-cell multiome libraries using the Chromium Next GEM Single-cell Multiome ATAC ^+^ Gene Expression kit (10X Genomics; 1000283) according to manufacturer’s instructions. Briefly, nuclei from monocytes were isolated based on 10X Genomics protocol recommendation (CG000365, Rev. D), loaded on Next GEM Chip J with a targeted nuclei recovery of 10,000 nuclei per donor and GEMs were generated using a 10X Chromium X series controller (10X Genomics; 00013-03) in-house. Library generation and evaluation also performed in house using a Tapestation 4200 system (Agilent). CD14^+^ monocytes were prepared and used fresh to avoid freeze-thaw cycles between isolation and nuclei extraction.

### Western blotting

Protein extracts from sorted monocytes from each donor were prepared in 1x CHAPS Lysis buffer (Millipore-Sigma, S7705) containing protease and phosphatase inhibitor cocktail (Thermo Fisher Scientific, 78440). Protein concentration was measured using the Qubit Protein Broad Range (BR) Assay Kit (Thermo Fisher Scientific, Q33212,) on a Qubit 4 Fluorometer according to manufacturer’s instructions. A total of 5 μg of whole-cell lysates were resolved on 4% –12% Bis-Tris Plus precast gels (Thermo Fisher Scientific, NW4125) in 1X MOPS SDS running buffer (Thermo Fisher Scientific), and proteins were transferred to nitrocellulose membranes via the iBlot 3 Western blot transfer system according to manufacturer’s instructions. Total protein staining in post transfer membranes was done using No-Stain Protein Labeling Reagent (Thermo Fisher Scientific, A44449) according to manufacturer’s instructions. Membranes were blocked in 5% non-fat dry milk in 1x TBST (150 mM NaCl, 10 mM Tris–HCl, pH 8.0, 0.05% Tween 20) at room temperature for 1 h, then incubated with primary antibodies for 2 h at room temperature or at 4°C overnight with gentle agitation. Membranes were washed three times in 1 × TBST for 10 min with shaking, then incubated with secondary antibodies for 1 h shaking, washed three times in 1 × TBST for 10 min with shaking. All blots were incubated with SuperSignal™ West Atto Ultimate Sensitivity Substrate (Thermo Scientific, A38556) and. The following primary antibodies and dilutions were used: anti-p21 Waf1/Cip1 (12D1) (rabbit monoclonal, Cell Signaling Technology, 2947), anti-p27 Kip1 (D69C12) (rabbit monoclonal, Cell Signaling Technology, 3686), anti-53BP1 (rabbit polyclonal, Novus Biologicals, NB-100-304), anti-cJun (H-79) (rabbit polyclonal, Santa Cruz Biotechnology, sc-1694), anti-mouse-HRP (Cell Signaling Technology, 7076), anti-rabbit-HRP (Cell Signaling Technology, 7074).

### Preprocessing of bulk high-throughput sequencing data

Paired-end reads were processed aligned to the GRCh38.d1.v1 version of the human genome using bowtie using the local mode (RNA-seq) or with the following parameters for ATAC-seq: –N 0 –-no-mixed –-no-discordant –-maxins 2000 –x –-trim3. Adapters were removed using cutadapt. PCR and optical duplicates were removed were removed with PicardTools. Enriched regions ATAC-seq were identified using MACS using a relaxed q-value threshold of 0.01. Master peak sets were constructed using a custom bedops script that merges common peaks between samples as we previously described^58^.

### Self-organizing maps (SOMs) and correlation analysis

SOM portraits were generated using oposSOM with default parameters, using the normalized and debatched (donor for SA-βGal; batch for age) reads. Metagenes were visualized in a 60 x 60 grid of rectangular topology, wherein expression portraits are projected by metagene distance matrix similarity using a logarithmic fold-change scale. Correlation analysis and associated visualizations were performed simultaneously during the SOM run.

### Differential expression

Differentially expressed genes (DEGs) were using DESeq2. Raw reads per exon, using the GRCh38.107 genome model, were quantified using the summarizeOverlaps package and peaks/genes with at least 10 reads in at least 10 libraries were kept. Correction of batch effects was performed with limma using the nucleic purification date (for age DEGs) or the donor (for SA-βGal DEGs) as the surrogate variables. Data transformation (regularized-log transformation), exploratory visualization (PCA and hierarchical clustering) and differential expression analysis were performed with DESeq2 using the default parameters and base R functions. We focused on highly significant DEGs by using an adjusted p-value filter of 0.05. Validation of differential expression analysis was performed by plotting the raw counts of a subset of senescence and age-associated DEGs as shown in the main text.

### Weighted gene co-expression network analysis (WGCNA)

DEGs identified with DESeq2 were used as input for unsupervised clustering using WGCNA. We used the “signed” option with default parameters except for the soft thresholding power. The minimum size for the DEG modules was set to 200 features for the initial set of modules, which were then merged by a dissimilarity threshold of 0.3. WGCNA modules were functionally profiled using clusterProfiler using the Molecular Signatures Database Hallmark gene sets. Statistical significance was calculated by a hypergeometric test with a cut-off of an adjusted p-value of ≤ 0.1 with Benjamini–Hochberg correction. Integration of DARs with DEGs was achieved by annotation of DARs to the nearest gene using ChIPSeeker79 followed by filtering against the list of DEGs.

### Integration of bulk RNA-seq and ATAC-seq data

To evaluate chromatin remodeling near transcriptional changes during monocyte aging and senescence, DEGs in WGCNA modules were integrated with ATAC-seq peak data. Gene symbols were mapped to Entrez IDs, and Transcription Start Sites (TSS) were defined using the TxDb.Hsapiens.UCSC.hg38.knownGene reference. Cis-regulatory genomic windows^+/−^ 500kb from DEG TSS were constructed via GenomicRanges. ATAC-seq peaks overlapping these windows were isolated using subsetByOverlaps, and strand-aware peak-center-to-TSS distances were computed. To resolve many-to-one peak-to-gene mappings, peak-level accessibility log2 fold change values were collapsed to a gene-level arithmetic mean. These aggregated ATAC metrics were merged with paired RNA-seq log2 fold change values for both age and senescence DEGs. The transcriptional and chromatin coupling efficiency was statistically quantified using a two-sided, non-parametric Spearman rank correlation test (cor.test), with comparisons being not significant. Modality concordance was visually assessed using ggplot2 scatter plots featuring dual marginal rug plots to highlight distribution densities.

### Transcription factor footprinting

Footprinting, TF motif enrichment, and differential binding activity were performed with HINT-ATAC using the JASPAR position weight matrix database for vertebrate TFs on merged ATAC-seq datasets, focusing on cis-regulatory regions^+/−^ 500 kb from SA-βGal– and age-specific DEGs. TF co-binding was assessed across ≤500bp bins of differentially accessible chromatin using a hypergeometric test for each pairwise co-binding possibility for each condition (SA-βGal levels and age, respectively) and adjusted p-values for multiple testing using Bonferroni correction. Differential activity was determined using HINT-Differential using default parameters. Visualizations were generated using ComplexHeatmap and pheatmap with annotated DEG status from the bulk RNA-seq differential analysis per TF.

### Transcription factor networks and network metrics

We built TF hierarchical networks essentially as we described^42^, except that we focused on the SA-βGal and age cis-regulatory regions as described above. Each identified TF is represented as a node, connected by directed edges representing co-binding events along the chromatin and during the transition from SA-βGal-low to SA-βGal-med to SA-βGal-high or young to old. An edge with TF A as the source and TF B as the target indicates that TF A binds to at least 15% of the same chromatin regions as TF B. For each TF, we computed their normalized total number of bound regions and dynamicity and represented those properties in the networks as node size and node color, respectively. Each edge is associated with a weight in the interval [0, 1], representing the fraction of TF B binding instances previously occupied by TF A. We filtered the networks for edges with a weight higher than 0.10 and performed a transitive reduction to simplify the obtained networks while keeping essential topological features. Nodes included in the same strongly connected component (SCC), i.e., connected both by incoming and outgoing paths, were merged into a single node. We represented the identified transcriptomic module distribution for TFs included in the same SCC as Doughnut Charts. Given the complexity of the networks, we further processed the network table by generating heatmaps of the scaled values for dynamicity, bound regions, out-degree and prestimulation for all TFs across the three network layers, for both senescence and age.

### Preprocessing of single-cell multiome data

Raw paired-end reads from multiome experiments (7 donors; 4 young, 3 old) were preprocessed using the 10X cell-ranger-arc pipeline. Initial cell calling was conducted via EmptyDropsMultiome^79^, implementing a batch-aware rescue workflow: high-noise samples (S1–S3) were filtered strictly using a false discovery rate boundary (FDR < 0.001), while clean samples (S4–S7) utilized an additional conditional inflection-point rescue mapping (barcodeRanks) to recover viable cell barcodes from ambient background signatures. Multiomic barcodes passing selection were aligned to valid human chromosomes and built into Seurat objects containing paired RNA and Chromatin Assays annotated with EnsDb.Hsapiens.v86. The RNA modality was normalized using SCTransform, its dimensionality reduced using Principal Component Analysis (PCA), and clustered using a shared nearest neighbor (SNN) graph with UMAP visualization. Cellular identities were assigned by mapping query profiles to a multimodal PBMC reference^60^, discarding low-confidence predictions with a maximum level 2 prediction score < 0.5. For the ATAC modality, data underwent Term Frequency-Inverse Document Frequency (TF-IDF) normalization, feature selection, and Latent Semantic Indexing (LSI) reduction using Singular Value Decomposition (components 2–30) as described by the Stuart lab. Joint QC metrics were programmatically calculated across both modalities, including mitochondrial, ribosomal, non-coding nuclear, transcription start site (TSS) enrichment, nucleosome banding signals, and genomic blacklist fractions. Final high-quality cell matrices were isolated using a joint-filtering lattice: nFeature_RNA (100–25,000), nCount_RNA (500–7,000), percent.mt <= 65%, TSS.enrichment >= 0.8, nucleosome_signal <= 2, and nCount_ATAC >= 1,000. This permissive mitochondrial filter, when cross-validated by strong, intact nuclear chromatin structural metrics (TSS and nucleosome signal), aligns with recent data-driven quality control literature^80,81^ demonstrating that standard, low-threshold mitochondrial cutoffs selectively destroy highly viable, metabolically active myeloid lineages.

### scRNA-seq data analysis

For single-cell (sc) RNA-seq data, filtered multiome single-cell objects (S1–S7) were merged, and sample-level metadata for chronological age and biological sex were appended. Global RNA profiles were normalized using SCTransform, followed by initial Principal Component Analysis (PCA) and UMAP embedding. To harmonize sample-specific batch variations while retaining biological differences, data integration across sample layers was executed via Canonical Correlation Analysis (CCAIntegration). Shared Nearest Neighbor (SNN) graph-based clustering (resolution = 0.6) and final UMAP dimensional reduction were subsequently performed on the integrated co-embedding space. To characterize age-associated transcriptional drift, cell-type-specific differential expression analysis was carried out on single-cells using FindMarkers within the isolated CD14^+^ monocyte subset. Shared signatures between single-cell and bulk profiles were cross-referenced against baseline age and senescence modules using UpSetR intersection frameworks. Statistically significant differentially expressed genes (adjusted p-value < 0.05, absolute log2 fold-change > 0.25) were highlighted via EnhancedVolcano plots and row-scaled hierarchical heatmaps (top 100 age DEGs). Functional over-representation analysis was using clusterProfiler, utilizing MSigDB Hallmark signature sets. Cluster-specific compositional shifts between the young and old cohorts were calculated as cell fractions, and variations in cluster membership were statistically evaluated for significance via pairwise Fisher’s exact tests. To quantify global module activity at single-cell resolution, UCell scoring was applied to evaluate senescence– and age-associated transcriptional programs (bulk senescence [SenSig] and age [AgeSig] as well as young– and old-specific DEGs from single-cell data [scYoung and scold, respectively]), projected onto the integrated UMAPs and their score distribution profiles rendered using kernel density frequency distributions.

### scATAC-seq data analysis

Data were normalized using Term Frequency-Inverse Document Frequency (TF-IDF) normalization and subjected to top feature selection. Dimensionality reduction was performed via Singular Value Decomposition to calculate Latent Semantic Indexing (LSI) components. Initial clustering and UMAP visualizations were executed using LSI components 2–30 to bypass depth-correlated variation, as per the Stuart lab guidelines. To resolve donor-specific batch effects, individual sample objects were split, integrated via reciprocal LSI (rlsi) space anchoring and integration was assessed by graph-based clustering and UMAP visualization. Differentially accessible regions (DARs) between young and old cohorts were determined by logistic regression modeling adjusted for ATAC sequencing depth (nCount_ATAC) as a latent variable, although most identified DARs were not statistically significant. To correlate to the gene expression output, DARs were mapped to their nearest upstream or downstream gene loci using GenomicRanges and the hg38 genome annotation. Peak-to-gene mappings were filtered to capture peak elements demonstrating the maximum log2 fold-change per gene. These summarized chromatin remodeling events were merged with the scRNA-seq differential expression matrix, categorized into modular significance quadrants, and statistically evaluated using a linear regression model and Spearman rank correlation test. For TF dynamics, overrepresented sequence motifs within DAR footprints were identified, and continuous single-cell transcription factor activity was estimated using chromVAR. Differentially active TFs between the young and old CD14^+^ monocyte subsets were isolated utilizing row means to evaluate the directional shift (avg_diff). The top 100 variable TF deviations were highlighted using row-scaled complete-linkage heatmaps split by age, sex, and cluster identifiers. As a complementary approach, we performed HINT footprinting and HINT-Differential using sample-specific pseudobulked scATAC-profiles (bed file export) using the scATAC master peakset as input, essentially as performed for bulk data and with annotated DEG status from the scRNA differential analysis per TF.

### Integrative pseudotime, RNA velocity, single-cell TF network dynamics and chromatin structural reorganization

To map the gene-regulatory transitions monocyte aging and senescence, we constructed a unified trajectory model using Monocle3. Combined cellular objects were transformed into a CellDataSet using the active SCT assay data layer and the trajectory constrained by projecting the Monocle3 trajectory onto the Seurat UMAP embeddings. Trajectory partitions and branches were resolved via cluster_cells and learn_graph with these parameters: use_partition = TRUE, close_loop = FALSE. Cells were ordered along a continuous pseudotime vector anchored by a root cell defined as the ‘youngest’ CD14-positive monocyte (minimum UMAP-1 value). Trajectory paths and path directionality were validated independently within a SingleCellExperiment environment using the slingshot package. The transition through senescence was validated by performing RNA velocity analysis. Briefly, spliced and unspliced transcript counts were generated from the Cell Ranger ARC output using Velocyto, and the resulting loom files from all samples were combined into a single AnnData object. The velocity dataset was matched to the integrated Seurat CCA analysis to retain the corresponding cell identities and UMAP coordinates. Using Scanpy, the data were preprocessed, and a neighborhood graph was constructed, after which scVelo was used to estimate RNA velocities with the stochastic model and compute a cell-to-cell transition graph. The inferred velocity vectors were projected onto the integrated CCA UMAP as velocity streamlines, providing a dynamic visualization of the predicted transcriptional trajectories and state transitions among monocyte populations. For global transcriptional entropy along the Monocle3 trajectory, a custom vector loop to measure absolute Shannon entropy metrics (H = –sum(p * log2(p))), extracting rolling generalized additive models (GAM) via mgcv to model critical state transitions was used. To evaluate pseudo-temporal kinetics systematically, the trajectory was divided into five bins based on defined pseudotime cutoffs: “Start”, “Pre-Bridge”, “Bridge” (the transitional senescence state), “Post-Bridge”, and “End”. To monitor senescence– and age-associated global chromatin structural rearrangements, the sub-nucleosomal and mono-nucleosomal fragmentation rates were binned dynamically utilizing FragmentHistogram and GetFragmentData across the hg38 reference assembly, mapping continuous structural metrics and total fragment abundance levels across specific promoter (within 3kb from TSS) and distal (> 3kb from TSS) enhancer windows. Transcription factor motif architecture was mapped using vertebrate matrices from the JASPAR2022 repository via motifmatchr and modeled continuously across pseudo-temporal bins using single-cell chromVAR deviation scores. TF activity transitions were tracked by calculating absolute magnitude kinetics across the matrix, assigning TFs to early activating waves or late-acting effector phases based on an absolute maximum intensity peak threshold. Continuous variations were projected using row-smoothed rolling-average (k = 50) chronologically ordered heatmaps (pheatmap). To determine how these dynamic tracking shifts directly influenced localized transcriptional output, lineage-determining (SPI1, IRF1), and master senescence regulators (FOS::JUN, FOSL2, JUN; AP-1 family). To validate the increased activity of AP-1 TFs with the aging transcriptional output, 401 bp windows centered on candidate AP-1 dimer motifs were overlapped with broad 1 Mb (^+/−^ 500 kb) cis-regulatory windows flanking the aging DEGs. The resulting insertion matrix was extracted and depth-normalized per 10,000 fragments using nCount_ATAC to account for library size variation. Profiles were partitioned by donor age group and trajectory phase, smoothened via a 7-bp rolling mean to resolve Tn5 signal dips and custom scatter plots generated with ggplot2 and ggrepel. A Spearman correlation analysis was performed between AP-1 activity profiles and the expression of the aging DEGs.

### Integration with clinical sepsis scRNA-seq cohorts

To provide clinical relevance to the senescence and age-associated monocyte states from our single-cell dataset, we integrated it with monocyte scRNA-seq data from distinct clinical sepsis cohorts^66^. Raw expression count matrices and metadata were parsed using fread, converted to Seurat objects, aligned by common cellular barcodes, and preprocessed via SCTransform normalization, PCA, and UMAP reduction. Monocyte populations were isolated from the clinical sepsis matrix and merged with the aging monocyte dataset. Unique cell prefix identifiers were appended to prevent barcode collisions, the datasets integrated using CCA) and evaluated with UMAPs. Spatial affinity and state transitions were quantified on UMAP coordinates by identifying the nearest clinical neighbors (k = 10) using k-nearest neighbors searching, followed by two-dimensional kernel density estimation and Euclidean distance profiling from clinical centroids (MS1–MS4), with statistical differences evaluated via Wilcoxon rank-sum tests with Benjamini-Hochberg correction. Transcriptional signatures for aging, senescence, and monocyte subsets (AgeSig, SenSig, scYoung, scOld, c5) were projected at single-cell resolution UCell scoring. Signature expression dynamics across clinical cohorts and monocyte states (MS) were profiled using ridge density plots and evaluated for statistical shifts using post-hoc Dunn’s tests. Over-representation analysis of MS state markers was executed against MSigDB Hallmark gene sets.

### Integration with microarray and bulk RNA-seq clinical sepsis datasets and gene set enrichment analysis (GSEA)

Microarray (GSE46955) and bulk RNA-seq (GSE133822) data were downloaded from GEO and SRA, respectively. Data were preprocessed and normalized, condition-specific DEGs identified using limma (microarray) or DESeq2 (bulk RNA-seq) and subsequently clustered using WGCNA. Enrichment of our transcriptional signatures for aging, senescence, and monocyte subsets (AgeSig, SenSig, scYoung, scOld, c5) was evaluated using clusterProfiler and enrichment calculated as described in the respective figure legends. For GSEA, we tested the prognostic potential of single-cell RNA-seq derived signatures (scOld and c5) in identifying septic and critically ill patients relative to healthy individuals from their global transcriptome. Briefly, we used the GSEA GUI version 4.2.2. Probe sets were collapsed to the gene level using the correlation-based approach. The correlation of probe sets representing the same gene was computed to decide whether to average probe sets (c > 0.2) or to use the probe set with the highest average expression across samples (c ≤ 0.2). Probe sets without known annotation were removed. The signal-to-noise ratio (μA – μB)/(σA ^+^ σB) (μ represents the mean, σ the standard deviation) was used as a ranking metric and statistics based on 500 gene set permutations.

## Funding

This work was supported by an American Federation for Aging Research and Glenn Foundation Award for Junior Faculty and a grant from the National Institute of General Medical Sciences of the National Institutes of Health (R35GM155447) to R.I.M.-Z. and grants R21AG067368 and R21AG067368-02S1 from the National Institute on Aging of the National Institutes of Health to P.F.-B.

## Contributions

R.I.M.-Z. and T.V. designed the study and prepared the manuscript. P.F.-B. and E.A. facilitated donor recruitment, blood collection and obtained flow cytometric SA-βGal data from PBMCs. R.I.M.-Z., T.V., P.S.T and L.G.-M. isolated senescent and non-senescent monocytes and performed high-throughput sequencing and flow cytometry experiments. P.S.T. and L.G.-M. analyzed flow cytometry data. T.V. performed single-cell multiome experiments. R.I.M.-Z. and TV analyzed high-throughput sequencing data. M.K. developed computational pipelines.

## Figures and Figure Legends

**Extended Data Figure 1.**
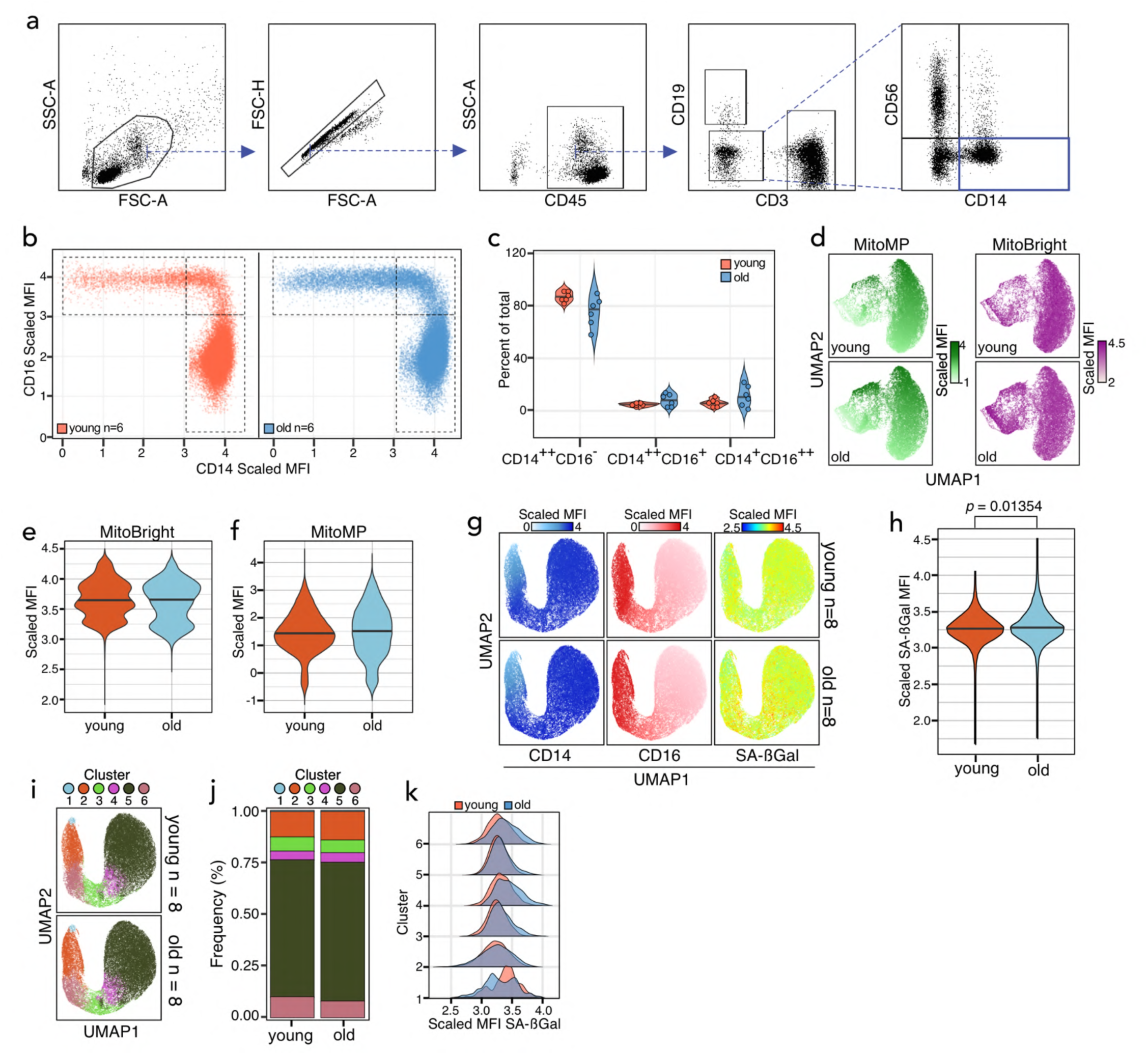
Increased SA-βGal activity in monocytes of older humans. **a.** Gating strategy for the analysis in Figure 1A. CD14-positive monocytes are highlighted by the teal box. **b.** Scatter plot showing the scaled MFI for CD14 and CD16 in younger and older donors. The gates used for quantification of monocyte subsets are shown. **c.** Violin plots showing the per-donor distribution of classical (CD14^++^CD16^-^), intermediate (CD14^++^CD16^+^) and non-classical (CD14^+^CD16^++^) monocytes. No statistical significance at the donor level was found using a Wilcoxon rank-sum test. **d.** UMAPs showing the scaled MFI for MitoMP and MitoBright in the younger and older donors from a. **e, f.** Violin plots for the scaled MFI of MitoMP and MitoBright in younger and older donors from A. No statistical significance was found using a Wilcoxon rank-sum test. **g.** Aggregate UMAPs for CD14, CD16 and SA-βGal scaled MFI. **h.** Violin plot showing the distribution shifts of scaled SA-βGal MFI values between younger and older donors. *P*-value was determined using a Wilcoxon rank-sum test from a sample of 1,000 random cells per age group from a total dataset of 60,000 monocytes (5,000 cells per donor per age group). **i.** UMAP showing FlowSOM clusters determined from input variables FSC-A, SSC-A, CD14, CD16, and SA-βGal from a cohort of 8 young and 8 older donors. **j.** Frequency histogram showing the FlowSOM cluster distribution shifts between younger and older donors. **k.** Density plots showing the shifts in distribution of scaled SA-βGal MFI values across FlowSOM clusters from h.

**Extended Data Figure 2.**
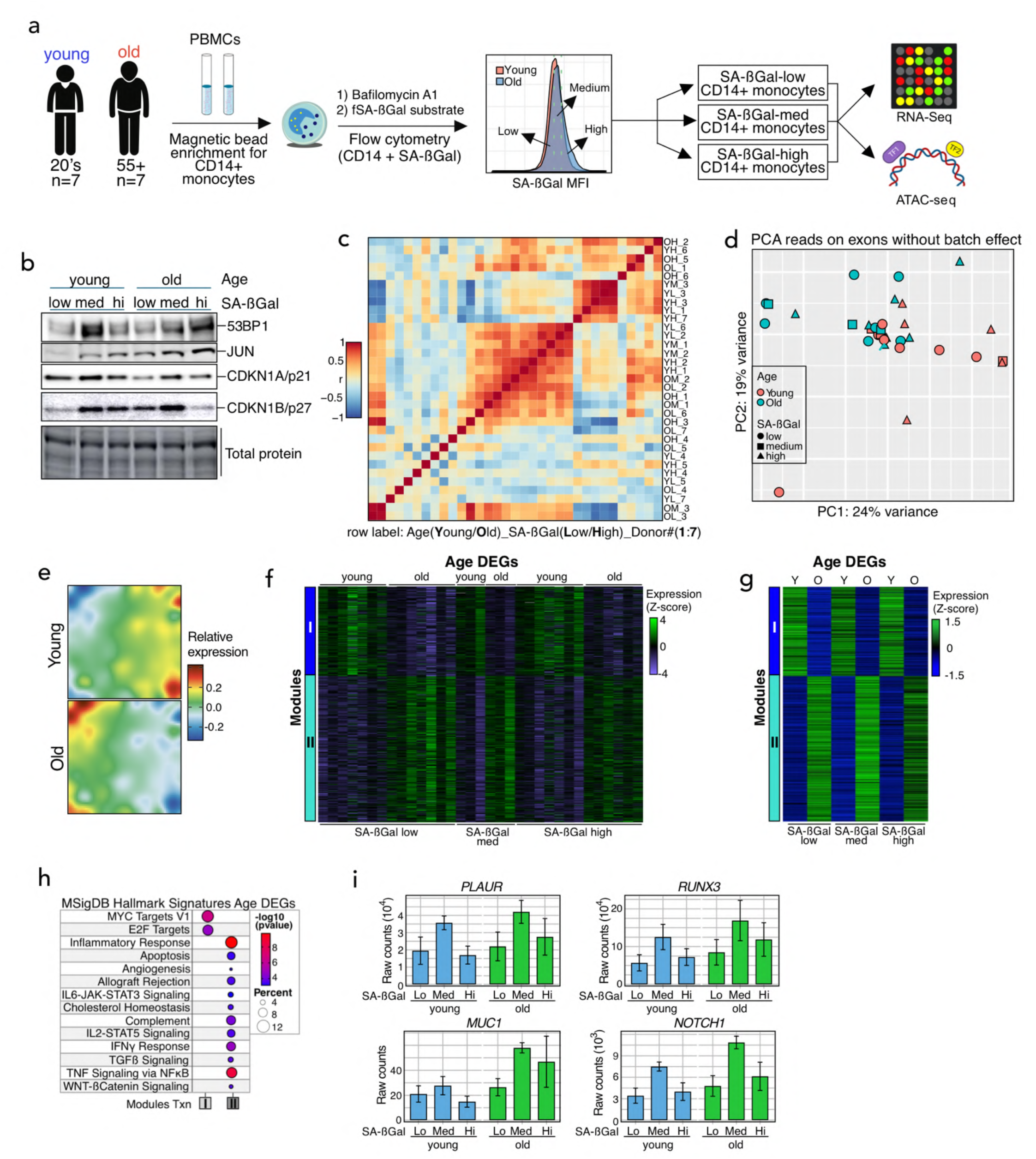
Distinct senescence and aging transcriptional programs in human monocytes. **a.** Schematic showing the strategy for the isolation and multiomic profiling of SA-βGal-low, –medium and –high CD14-positive monocytes from younger and older donors. **b.** Immunoblot characterization of the indicated senescence-associated molecular markers in SA-βGal-low, –medium and –high CD14-positive monocytes from a younger and an older donor. **c.** Correlation heatmap between the transcriptomes of SA-βGal-low, –medium and –high CD14^+^ monocytes of older and younger donors after correcting for batch variation. **d.** PCA projection plot of reads on exons of the indicated transcriptomes of SA-βGal-low, –medium and –high CD14-positive monocytes from older and younger donors after correcting for batch variation. **e.** Averaged SOMs of donor variation-corrected transcriptomes of CD14-positive monocytes of younger and older donors. **f, g.** Individual (f) and averaged (g) heatmaps of number-coded modules of DEGs in the transcriptomes of CD14-positive monocytes of the indicated younger and older donors. The scale bar indicates the row z-score of the rlog-transformed counts. **h.** Functional overrepresentation analysis map showing significant associations of the MSigDB hallmark gene sets for each module described in (f, g). Circle fill is color coded according to the false discovery rate (FDR)-corrected *p*-value from a hypergeometric distribution test. Circle size is proportional to the percentage of genes in each MSigDB gene set. **i.** Bar plots showing the average raw counts of a subset of genes displaying increased expression in an age-dependent manner in the transcriptomes of younger and older donors. Data from the following number of donors: SA-βGal-low, n=7; SA-βGal-med, n=3; SA-βGal-high, n=7; young, n=7; old, n=7 (A-I).

**Extended Data Figure 3.**
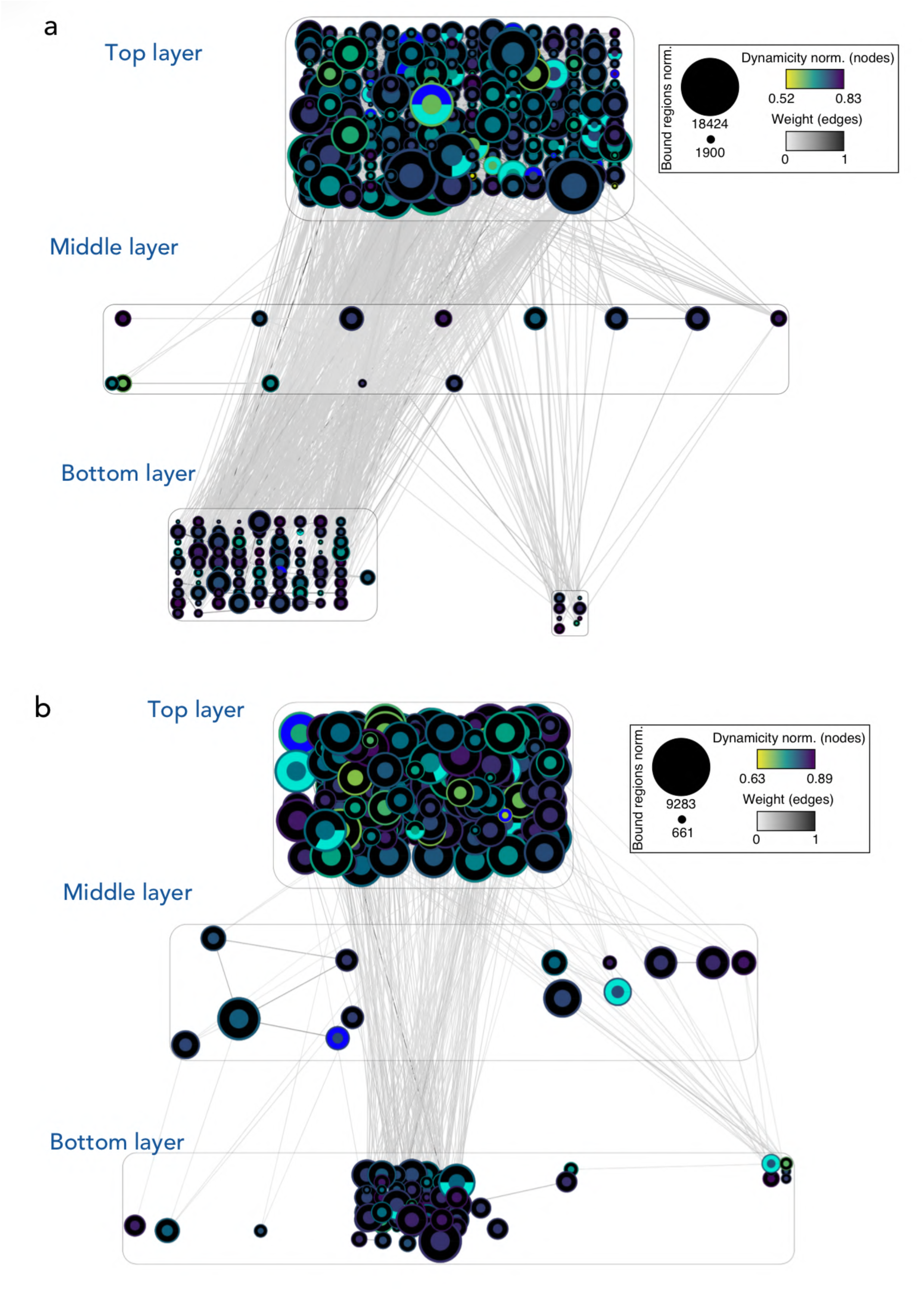
High complexity of the TF networks controlling monocyte senescence and aging transcriptional programs. **a, b.** Graphical representations of the full TF networks at the accessible chromatin (^+/−^ 500kb) of SA-βGal– and age-associated DEGs (a, b). TFs (nodes) are represented as circles. Oriented edges (arrows) connecting nodes indicate that at least 10% of the regions bound by a given TF in the bottom and core layers were bound by the interacting TF in the core and top layers, respectively, at the same or previous time points. The fill color of the node’s inner circle is based on the normalized dynamicity of TFs. The fill color of the outer ring indicates whether the TF is constitutively expressed (black) or belongs to a transcriptomic module (Figures 2 and Extended Data Figure 2). The node’s size is proportional to the bound regions by a given TF. Each network has three layers: (1) the top layer with no incoming edges, (2) the core layer with incoming and outgoing edges, and (3) the bottom layer with no outgoing edges. The inset denotes the bound regions scale, prestimulation and directional overlap of the TF interactions. For ease of visualization, edges have been bundled. Networks were constructed from chromatin accessibility data from the following donors: SA-βGal-low, n=7; SA-βGal-med, n=3; SA-βGal-high, n=7; young, n=7; old, n=7 (a, b).

**Extended Data Figure 4.**
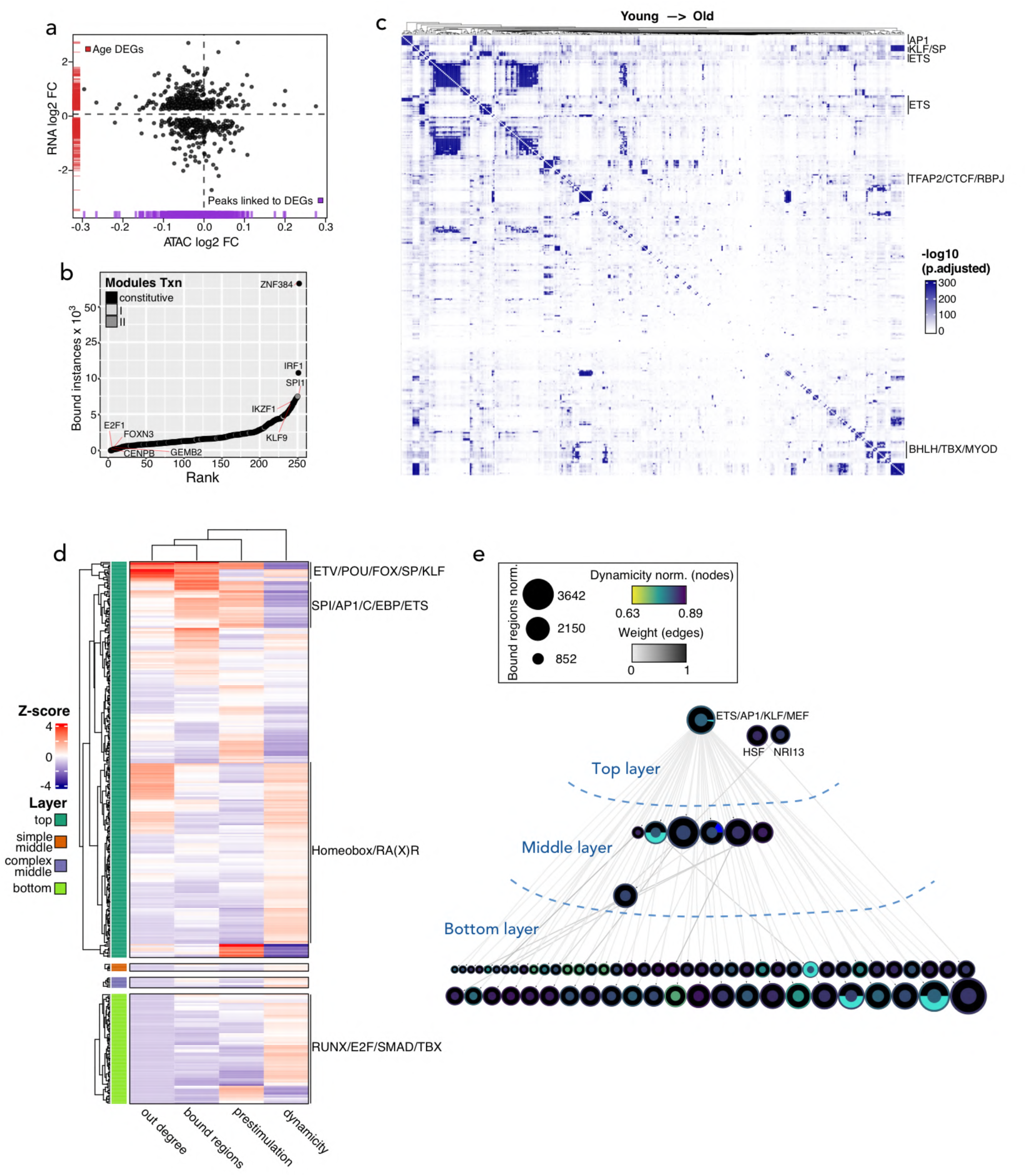
Widespread TF network reorganization during the transitions to senescence and aged monocyte states. **a.** Correlation plot of the RNA log2-fold change of age-associated DEGs and the log2-fold change in the accessibility of their linked peaks (^+/−^ 500kb). Each black dot represents a DEG linked to at least one chromatin accessibility peak, with its x-axis position reflecting the mean log2-fold change across all peaks linked to that gene. Red ticks on the y-axis represent the RNA log2-fold change distribution of all age DEGs. Blue ticks on the x-axis represent the distribution of mean chromatin accessibility log2-fold changes for each SA-βGal DEG. **b.** Rank plot showing the summed binding instances of TFs at age DEG-associated chromatin accessibility peaks (^+/−^ 500kb). **c.** Co-binding matrices of TF interactions (500 bp resolution; Ward’s criterion for clustering) age DEG-associated peaks. The corresponding q values of the interactions are projected onto the clustering and represented in a color scale defined by their significance using a hypergeometric distribution test. **d.** Heatmap showing the variability of TFs in each layer of the age-specific networks across dynamicity, number of bound regions, interactions (out degree) and prestimulation. The scale bar indicates the row Z score of the metric value of each TF motif in the respective network layer. **e.** Simplified TF networks at the accessible chromatin (^+/−^ 500kb) of age-associated DEGs. Nodes represent strongly connected components to facilitate visualization. The fill color of the node’s inner circle is based on the normalized dynamicity (prestimulation) of TFs. The fill color of the outer ring indicates whether the TF is constitutively expressed (black) or belongs to a transcriptomic module as described in Figure 2f. The node’s size is proportional to the bound regions by a given TF(s). Each network has three layers: (1) the top layer with no incoming edges, (2) the core layer with incoming and outgoing edges, and (3) the bottom layer with no outgoing edges. The inset denotes the bound regions scale, prestimulation and directional overlap of the TF interactions. The network was constructed from chromatin accessibility data from 7 young and 7 old donors.

**Extended Data Figure 5.**
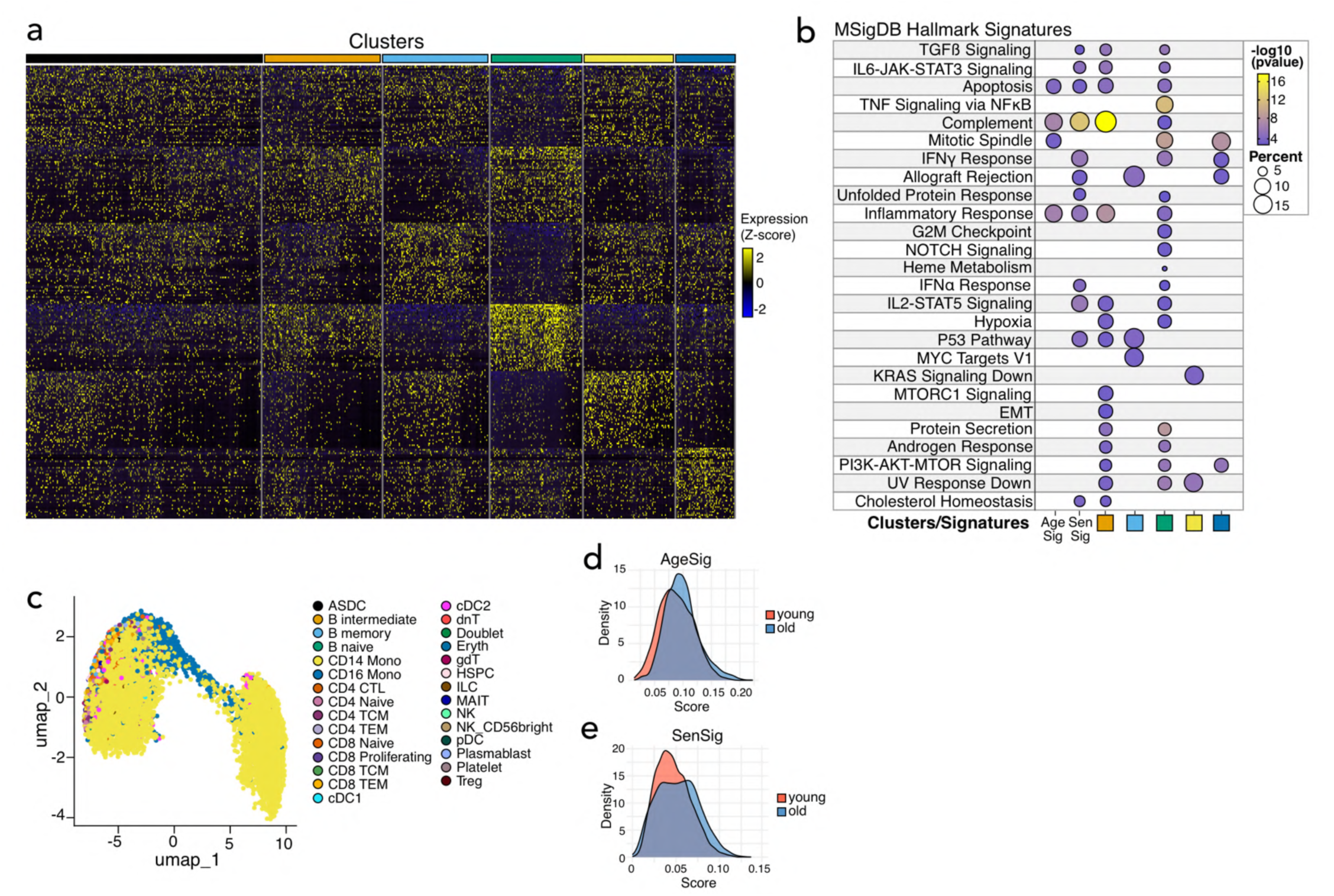
A transitional senescence state leads to an age-associated inflammatory monocyte state. **a.** Heatmap showing the single-cell expression of the top 50 cluster-specific DEGs identified in Figure 5c. **b.** Functional overrepresentation analysis map showing significant associations of the MSigDB hallmark gene sets for each cluster-specific DEG set from a. Circle fill is color coded according to the false discovery rate (FDR)-corrected *p*-value from a hypergeometric distribution test. Circle size is proportional to the percentage of genes in each MSigDB gene set. **c.** UMAP showing the cell type annotation using label transfer from the Azimuth PBMC reference (prediction score >= 0.5). The overwhelming majority of cells are labelled as CD14^+^ and CD16^+^ monocytes. **d, e.** Overlapping density plots displaying the single-cell UCell enrichment scores for the age (AgeSig) and senescence (SenSig) gene signatures derived from bulk RNA-seq sets from Figure 2 and Extended Data Figure 2. Note the increased representation of both signatures in single CD14^+^ monocytes from older donors.

**Extended Data Figure 6.**
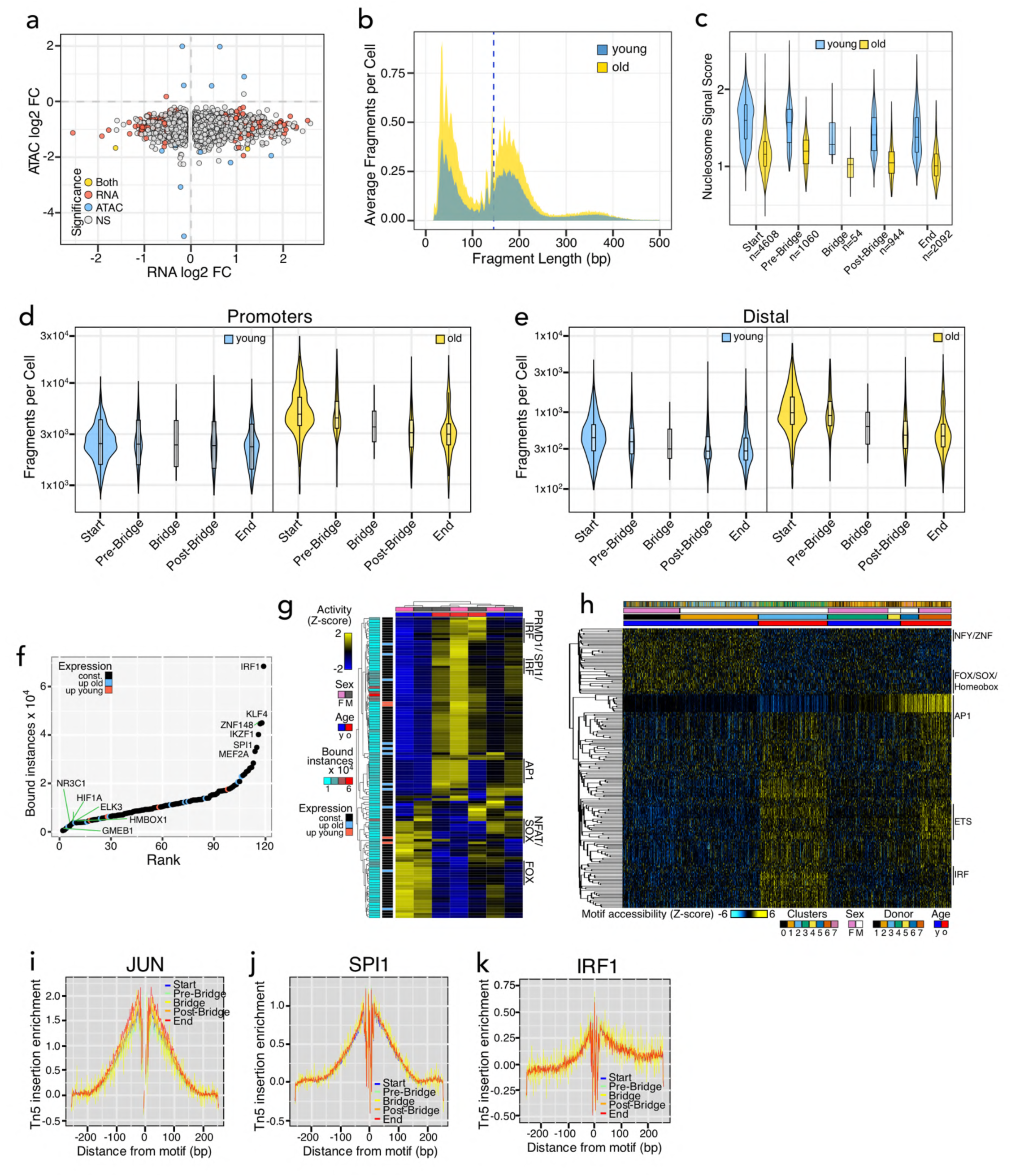
A transitional senescence state leads to an age-associated inflammatory monocyte state. **a.** Correlation plot of the single-cell RNA log2-fold change of age-associated DEGs and the single-cell log2-fold change of their nearest chromatin accessibility peaks. Each dot represents a DEG linked to at least one chromatin accessibility peak. Statistical significance was calculated using a two-sided Wilcoxon Rank Sum test for adjusted *p*-value < 0-05 and | log2FC > 0.35 | for either or both modalities. **b.** Fragment length distribution across a 10-megabase region of chromosome 1 (1-10,000,000). **c.** Violin plots showing the nucleosome signal score per age group along pseudotime. The number of cells per pseudotime bin is shown on the x-axis. **d, e.** Violin plots showing the fragments per cell at promoters (within 3kb from the TSS; d) and distal regions (^+/−^ 3kb from the TSS; e) per age group. **f.** Rank plot showing the summed binding instances of TFs at peaks with sample-specific differential binding activity on pseudobulked chromatin accessibility. **g.** Heatmap of sample-specific differential TF binding activity at accessible chromatin of monocytes. Chromatin accessibility data per sample was pseudobulked and differential binding activity was determined using RGT-HINT. The left annotation heatmaps show the number of bound instances per TF (cyan to red) and their gene expression (TXN) category as determined in d. The column annotation refers to the sex and age of each sample. A subset of TF families showing age-specific activity is highlighted. The scale bar indicates the row Z score of the activity of each TF motif in the dataset. **h.** Heatmap showing the motif accessibility dynamics of the top 200 variable TFs between age groups at single-cell resolution. The top annotations are defined at the bottom of the heatmap. A subset of TF families with age-specific binding behavior is indicated to the right of the heatmap. **i-k.** Genome-wide footprinting metaprofiles of JUN (i), SPI1/PU.1 (j) and IRF1 (k).

**Extended Data Figure 7.**
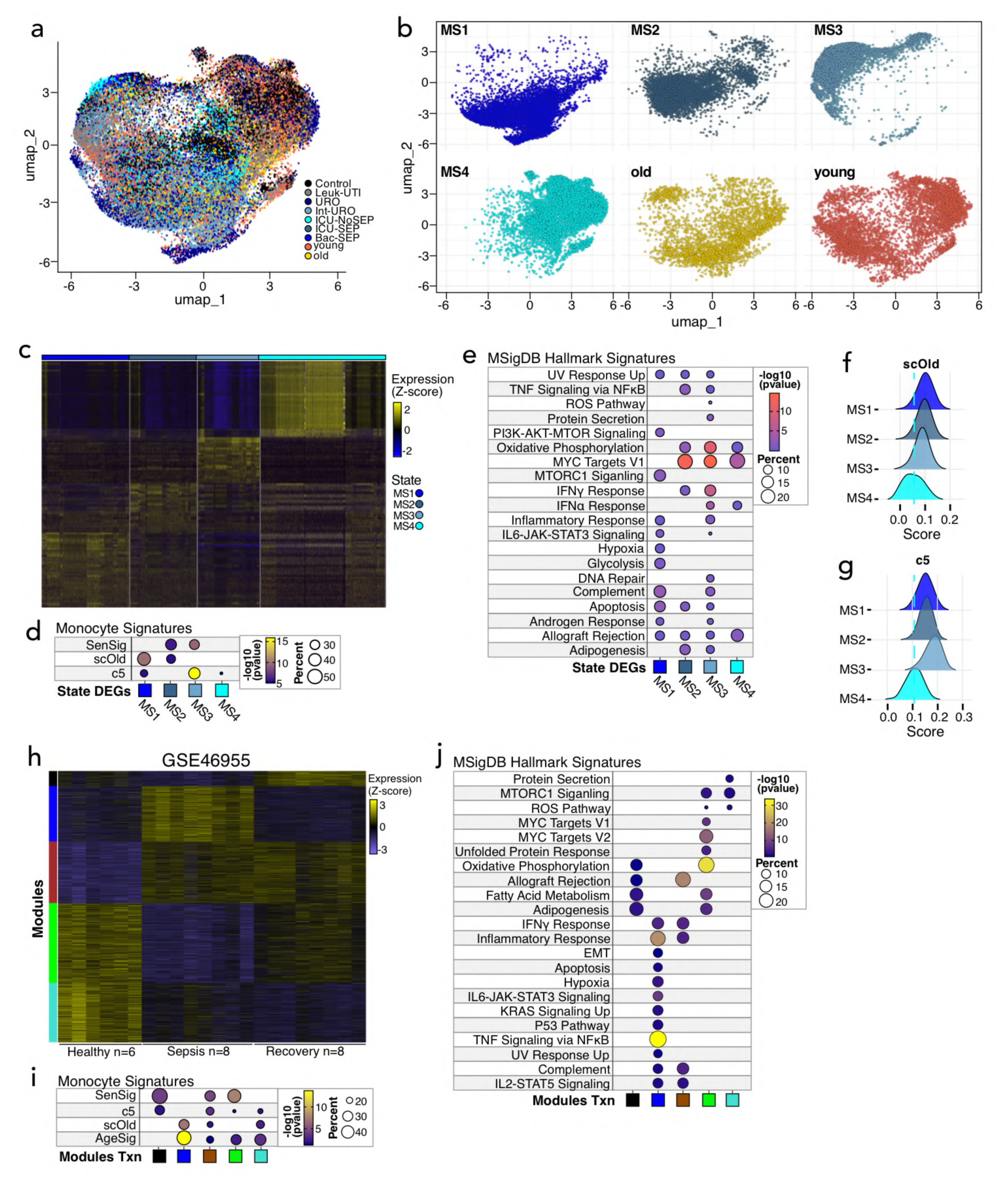
Accumulation of septic monocytes in aging humans. **a.** Overlaid integrated UMAP showing the spatial distribution of young and old monocytes (this study) relative to control and septic monocytes from the indicated sepsis cohorts from reference^66^. **b**. Split integrated UMAPs showing the spatial distribution of monocyte states from^66^ and young and old monocytes. **c**. Heatmap showing the single-cell expression of the top 50 monocyte state-specific DEGs. **d, e.** Functional overrepresentation analysis map showing significant associations of the indicated age and senescence signatures (d) and the MSigDB hallmark gene sets (e) for each monocyte state from c. **f, g.** Ridge plots showing the shift of the scOld and c5 gene signature scores across control and septic monocytes from reference^66^. The dashed cyan indicate the median score of each gene signature in MS4 monocytes. Statistical significance of gene signature score shifts in MS1-3 states relative to MS4 was calculated using post-hoc Dunn tests. *P*.adj < 2e^-16^ for all comparisons. **h.** Individual heatmaps of color-coded modules of DEGs in the transcriptomes of monocytes of patients in the indicated conditions from dataset GSE46955. **i, j**. Functional overrepresentation analysis map showing significant associations of the indicated age and senescence signatures (i) and the MSigDB hallmark gene sets (j) for each DEG module from h. Circle fill is color coded according to the false discovery rate (FDR)-corrected *p*-value from a hypergeometric distribution test. Circle size is proportional to the percentage of genes in each MSigDB gene set (d, e, i, j).

## Notes

### Competing Interest Statement

The authors have declared no competing interest.

## References

1 Campisi, J. Aging, cellular senescence, and cancer. Annu Rev Physiol 75, 685–705 (2013). 10.1146/annurev-physiol-030212-183653

2 Childs, B. G., Durik, M., Baker, D. J. & van Deursen, J. M. Cellular senescence in aging and age-related disease: from mechanisms to therapy. Nat Med 21, 1424–1435 (2015). 10.1038/nm.4000

3 Birch, J. & Gil, J. Senescence and the SASP: many therapeutic avenues. Genes Dev 34, 1565–1576 (2020). 10.1101/gad.343129.120

4 Coppe, J. P., Desprez, P. Y., Krtolica, A. & Campisi, J. The senescence-associated secretory phenotype: the dark side of tumor suppression. Annu Rev Pathol 5, 99–118 (2010). 10.1146/annurev-pathol-121808-102144

5 Majewska, J. & Krizhanovsky, V. Immune surveillance of senescent cells in aging and disease. Nat Aging 5, 1415–1424 (2025). 10.1038/s43587-025-00910-5

6 Jang, I. H., Niedernhofer, L. J., Robbins, P. D. & Camell, C. D. The ageing immune system as a driver of systemic ageing. Nat Rev Immunol 26, 489–506 (2026). 10.1038/s41577-026-01269-3

7 Jing, Y., et al. Aging is associated with a numerical and functional decline in plasmacytoid dendritic cells, whereas myeloid dendritic cells are relatively unaltered in human peripheral blood. Hum Immunol 70, 777–784 (2009). 10.1016/j.humimm.2009.07.005

8 Liu, Z. et al. Immunosenescence: molecular mechanisms and diseases. Signal Transduct Target Ther 8, 200 (2023). 10.1038/s41392-023-01451-2

9 Goronzy, J. J. & Weyand, C. M. Understanding immunosenescence to improve responses to vaccines. Nat Immunol 14, 428–436 (2013). 10.1038/ni.2588

10 Turano, P. S., et al. Age-independent and targetable transcription factor networks regulating CD8(+) T cell senescence in aging humans. Cell Rep 45, 116795 (2026). 10.1016/j.celrep.2025.116795

11 Martinez-Zamudio, R. I., et al. Senescence-associated beta-galactosidase reveals the abundance of senescent CD8+ T cells in aging humans. Aging Cell 20, e13344 (2021). 10.1111/acel.13344

12 Pereira, B. I., et al. Senescent cells evade immune clearance via HLA-E-mediated NK and CD8(+) T cell inhibition. Nat Commun 10, 2387 (2019). 10.1038/s41467-019-10335-5

13 Callender, L. A., et al. Human CD8(+) EMRA T cells display a senescence-associated secretory phenotype regulated by p38 MAPK. Aging Cell 17 (2018). 10.1111/acel.12675

14 Mognol, G. P., et al. Exhaustion-associated regulatory regions in CD8(+) tumor-infiltrating T cells. Proc Natl Acad Sci U S A 114, E2776–E2785 (2017). 10.1073/pnas.1620498114

15 Wherry, E. J., et al. Molecular signature of CD8+ T cell exhaustion during chronic viral infection. Immunity 27, 670–684 (2007). 10.1016/j.immuni.2007.09.006

16 Patel, A. A., et al. The fate and lifespan of human monocyte subsets in steady state and systemic inflammation. J Exp Med 214, 1913–1923 (2017). 10.1084/jem.20170355

17 Guilliams, M., Mildner, A. & Yona, S. Developmental and Functional Heterogeneity of Monocytes. Immunity 49, 595–613 (2018). 10.1016/j.immuni.2018.10.005

18 Geissmann, F., et al. Development of monocytes, macrophages, and dendritic cells. Science 327, 656– 661 (2010). 10.1126/science.1178331

19 Ziegler-Heitbrock, L., et al. Nomenclature of monocytes and dendritic cells in blood. Blood 116, e74–80 (2010). 10.1182/blood-2010-02-258558

20 Swirski, F. K., et al. Identification of splenic reservoir monocytes and their deployment to inflammatory sites. Science 325, 612–616 (2009). 10.1126/science.1175202

21 Scott, C. L., et al. Bone marrow-derived monocytes give rise to self-renewing and fully differentiated Kupffer cells. Nat Commun 7, 10321 (2016). 10.1038/ncomms10321

22 Chambers, E. S., et al. Recruitment of inflammatory monocytes by senescent fibroblasts inhibits antigen-specific tissue immunity during human aging. Nat Aging 1, 101–113 (2021). 10.1038/s43587-020-00010-6

23 Henderson, R. B., Hobbs, J. A., Mathies, M. & Hogg, N. Rapid recruitment of inflammatory monocytes is independent of neutrophil migration. Blood 102, 328–335 (2003). 10.1182/blood-2002-10-3228

24 Belge, K. U., et al. The proinflammatory CD14+CD16+DR++ monocytes are a major source of TNF. J Immunol 168, 3536–3542 (2002). 10.4049/jimmunol.168.7.3536

25 Tacke, F., et al. Immature monocytes acquire antigens from other cells in the bone marrow and present them to T cells after maturing in the periphery. J Exp Med 203, 583–597 (2006). 10.1084/jem.20052119

26 Wong, K. L., et al. Gene expression profiling reveals the defining features of the classical, intermediate, and nonclassical human monocyte subsets. Blood 118, e16–31 (2011). 10.1182/blood-2010-12-326355

27 Lee, J., et al. The MHC class II antigen presentation pathway in human monocytes differs by subset and is regulated by cytokines. PLoS One 12, e0183594 (2017). 10.1371/journal.pone.0183594

28 Cros, J., et al. Human CD14dim monocytes patrol and sense nucleic acids and viruses via TLR7 and TLR8 receptors. Immunity 33, 375–386 (2010). 10.1016/j.immuni.2010.08.012

29 Wallis, Z. K. & Williams, K. C. Monocytes in HIV and SIV Infection and Aging: Implications for Inflamm-Aging and Accelerated Aging. Viruses 14 (2022). 10.3390/v14020409

30 De Maeyer, R. P. H., et al. Age-Associated Inflammatory Monocytes Are Increased in Menopausal Females and Reversed by Hormone Replacement Therapy. Aging Cell 24, e70249 (2025). 10.1111/acel.70249

31 Costantini, A., et al. Age-related M1/M2 phenotype changes in circulating monocytes from healthy/unhealthy individuals. Aging (Albany NY) 10, 1268–1280 (2018). 10.18632/aging.101465

32 Snodgrass, R. G., Jiang, X. & Stephensen, C. B. Monocyte subsets display age-dependent alterations at fasting and undergo non-age-dependent changes following consumption of a meal. Immun Ageing 19, 41 (2022). 10.1186/s12979-022-00297-6

33 Hearps, A. C., et al. Aging is associated with chronic innate immune activation and dysregulation of monocyte phenotype and function. Aging Cell 11, 867–875 (2012). 10.1111/j.1474-9726.2012.00851.x

34 Rimpa, C. M., et al. Characterization of the Molecular Signature of Human Monocytes in Aging and Myelodysplastic Neoplasms. Eur J Immunol 55, e202451387 (2025). 10.1002/eji.202451387

35 Seidler, S., Zimmermann, H. W., Bartneck, M., Trautwein, C. & Tacke, F. Age-dependent alterations of monocyte subsets and monocyte-related chemokine pathways in healthy adults. BMC Immunol 11, 30 (2010). 10.1186/1471-2172-11-30

36 Metcalf, T. U., et al. Human Monocyte Subsets Are Transcriptionally and Functionally Altered in Aging in Response to Pattern Recognition Receptor Agonists. J Immunol 199, 1405–1417 (2017). 10.4049/jimmunol.1700148

37 Saare, M., et al. Monocytes present age-related changes in phospholipid concentration and decreased energy metabolism. Aging Cell 19, e13127 (2020). 10.1111/acel.13127

38 Pence, B. D. & Yarbro, J. R. Aging impairs mitochondrial respiratory capacity in classical monocytes. Exp Gerontol 108, 112–117 (2018). 10.1016/j.exger.2018.04.008

39 Ruiz, V. Y., et al. Single-cell analysis of CD14(+)CD16(+) monocytes identifies a subpopulation with an enhanced migratory and inflammatory phenotype. Front Immunol 16, 1475480 (2025). 10.3389/fimmu.2025.1475480

40 Olinger, B., et al. The secretome of senescent monocytes predicts age-related clinical outcomes in humans. Nat Aging 5, 1266–1279 (2025). 10.1038/s43587-025-00877-3

41 Lorente-Sorolla, C., et al. Inflammatory cytokines and organ dysfunction associate with the aberrant DNA methylome of monocytes in sepsis. Genome Med 11, 66 (2019). 10.1186/s13073-019-0674-2

42 Martinez-Zamudio, R. I. et al. AP-1 imprints a reversible transcriptional programme of senescent cells. Nat Cell Biol 22, 842–855 (2020). 10.1038/s41556-020-0529-5

43 Martinez-Zamudio, R. I., et al. Escape from oncogene-induced senescence is controlled by POU2F2 and memorized by chromatin scars. Cell Genom 3, 100293 (2023). 10.1016/j.xgen.2023.100293

44 Kaczorowski, K. J., et al. Continuous immunotypes describe human immune variation and predict diverse responses. Proc Natl Acad Sci U S A 114, E6097–E6106 (2017). 10.1073/pnas.1705065114

45 Kroger, C., et al. Unveiling the power of high-dimensional cytometry data with cyCONDOR. Nat Commun 15, 10702 (2024). 10.1038/s41467-024-55179-w

46 Yarbro, J. R. & Pence, B. D. Classical monocytes from older adults maintain capacity for metabolic compensation during glucose deprivation and lipopolysaccharide stimulation. Mech Ageing Dev 183, 111146 (2019). 10.1016/j.mad.2019.111146

47 Langfelder, P. & Horvath, S. WGCNA: an R package for weighted correlation network analysis. BMC Bioinformatics 9, 559 (2008). 10.1186/1471-2105-9-559

48 De Cecco, M., et al. L1 drives IFN in senescent cells and promotes age-associated inflammation. Nature 566, 73–78 (2019). 10.1038/s41586-018-0784-9

49 Yu, Q., et al. DNA-damage-induced type I interferon promotes senescence and inhibits stem cell function. Cell Rep 11, 785–797 (2015). 10.1016/j.celrep.2015.03.069

50 Patrick, R., et al. The activity of early-life gene regulatory elements is hijacked in aging through pervasive AP-1-linked chromatin opening. Cell Metab 36, 1858–1881 e1823 (2024). 10.1016/j.cmet.2024.06.006

51 Byrns, C. N., Saikumar, J. & Bonini, N. M. Glial AP1 is activated with aging and accelerated by traumatic brain injury. Nat Aging 1, 585–597 (2021). 10.1038/s43587-021-00072-0

52 Zhang, C., et al. ATF3 drives senescence by reconstructing accessible chromatin profiles. Aging Cell 20, e13315 (2021). 10.1111/acel.13315

53 Buenrostro, J. D., Giresi, P. G., Zaba, L. C., Chang, H. Y. & Greenleaf, W. J. Transposition of native chromatin for fast and sensitive epigenomic profiling of open chromatin, DNA-binding proteins and nucleosome position. Nat Methods 10, 1213–1218 (2013). 10.1038/nmeth.2688

54 Garber, M., et al. A high-throughput chromatin immunoprecipitation approach reveals principles of dynamic gene regulation in mammals. Mol Cell 47, 810–822 (2012). 10.1016/j.molcel.2012.07.030

55 Kurotaki, D., et al. Chromatin structure undergoes global and local reorganization during murine dendritic cell development and activation. Proc Natl Acad Sci U S A 119, e2207009119 (2022). 10.1073/pnas.2207009119

56 Natoli, G. Maintaining cell identity through global control of genomic organization. Immunity 33, 12–24 (2010). 10.1016/j.immuni.2010.07.006

47 Dickerson, K. M. et al. ZNF384 Fusion Oncoproteins Drive Lineage Aberrancy in Acute Leukemia. Blood Cancer Discov 3, 240–263 (2022). 10.1158/2643-3230.BCD-21-0163

58 Turano, P. S., et al. Age-independent and targetable transcription factor networks regulating CD8(+) T cell senescence in aging humans. Cell Rep 45, 116795 (2025). 10.1016/j.celrep.2025.116795

59 Tarjan, R. Depth-First Search and Linear Graph Algorithms. SIAM Journal on Computing 1, 146–160 (1972). 10.1137/0201010

60 Hao, Y., et al. Integrated analysis of multimodal single-cell data. Cell 184, 3573–3587 e3529 (2021). 10.1016/j.cell.2021.04.048

61 Cao, J., et al. The single-cell transcriptional landscape of mammalian organogenesis. Nature 566, 496– 502 (2019). 10.1038/s41586-019-0969-x

62 Bergen, V., Lange, M., Peidli, S., Wolf, F. A. & Theis, F. J. Generalizing RNA velocity to transient cell states through dynamical modeling. Nat Biotechnol 38, 1408–1414 (2020). 10.1038/s41587-020-0591-3

63 Schep, A. N., Wu, B., Buenrostro, J. D. & Greenleaf, W. J. chromVAR: inferring transcription-factor-associated accessibility from single-cell epigenomic data. Nat Methods 14, 975–978 (2017). 10.1038/nmeth.4401

64 van der Poll, T., van de Veerdonk, F. L., Scicluna, B. P. & Netea, M. G. The immunopathology of sepsis and potential therapeutic targets. Nat Rev Immunol 17, 407–420 (2017). 10.1038/nri.2017.36

65 Shaw, A. C., Goldstein, D. R. & Montgomery, R. R. Age-dependent dysregulation of innate immunity. Nat Rev Immunol 13, 875–887 (2013). 10.1038/nri3547

66 Reyes, M., et al. An immune-cell signature of bacterial sepsis. Nat Med 26, 333–340 (2020). 10.1038/s41591-020-0752-4

67 Shalova, I. N., et al. Human monocytes undergo functional re-programming during sepsis mediated by hypoxia-inducible factor-1alpha. Immunity 42, 484–498 (2015). 10.1016/j.immuni.2015.02.001

68 Washburn, M. L., et al. T Cell– and Monocyte-Specific RNA-Sequencing Analysis in Septic and Nonseptic Critically Ill Patients and in Patients with Cancer. J Immunol 203, 1897–1908 (2019). 10.4049/jimmunol.1900560

69 Pereira, B. I., et al. Sestrins induce natural killer function in senescent-like CD8(+) T cells. Nat Immunol 21, 684–694 (2020). 10.1038/s41590-020-0643-3

70 Elyahu, Y., et al. CD4 T cells acquire Eomesodermin to modulate cellular senescence and aging. Nat Aging 5, 1970–1982 (2025). 10.1038/s43587-025-00953-8

71 Chambers, E. S., et al. Recruitment of inflammatory monocytes by senescent fibroblasts inhibits antigen-specific tissue immunity during human aging. Nature Aging 1, 101–113 (2021). 10.1038/s43587-020-00010-6

72 Kapellos, T. S., et al. Human Monocyte Subsets and Phenotypes in Major Chronic Inflammatory Diseases. Front Immunol 10, 2035 (2019). 10.3389/fimmu.2019.02035

73 Gren, S. T., et al. A Single-Cell Gene-Expression Profile Reveals Inter-Cellular Heterogeneity within Human Monocyte Subsets. PLoS One 10, e0144351 (2015). 10.1371/journal.pone.0144351

74 Reitsema, R. D., Kumawat, A. K., Hesselink, B. C., van Baarle, D. & van Sleen, Y. Effects of ageing and frailty on circulating monocyte and dendritic cell subsets. NPJ Aging 10, 17 (2024). 10.1038/s41514-024-00144-6

75 Hanai, H., et al. Adsorptive depletion of elevated proinflammatory CD14+CD16+DR++ monocytes in patients with inflammatory bowel disease. Am J Gastroenterol 103, 1210–1216 (2008). 10.1111/j.1572-0241.2007.01714.x

76 Koch, S., Kucharzik, T., Heidemann, J., Nusrat, A. & Luegering, A. Investigating the role of proinflammatory CD16+ monocytes in the pathogenesis of inflammatory bowel disease. Clin Exp Immunol 161, 332–341 (2010). 10.1111/j.1365-2249.2010.04177.x

77 Yin, Q., et al. The CD14(++)CD16(+) monocyte subset is expanded and controls Th1 cell development in Graves’ disease. Clin Immunol 245, 109160 (2022). 10.1016/j.clim.2022.109160

78 Rossol, M., Kraus, S., Pierer, M., Baerwald, C. & Wagner, U. The CD14(bright) CD16+ monocyte subset is expanded in rheumatoid arthritis and promotes expansion of the Th17 cell population. Arthritis Rheum 64, 671–677 (2012). 10.1002/art.33418

79 Megas, S., Lorenzi, V. & Marioni, J. C. EmptyDropsMultiome discriminates real cells from background in single-cell multiomics assays. Genome Biol 25, 121 (2024). 10.1186/s13059-024-03259-x

80 Subramanian, A., Alperovich, M., Yang, Y. & Li, B. Biology-inspired data-driven quality control for scientific discovery in single-cell transcriptomics. Genome Biol 23, 267 (2022). 10.1186/s13059-022-02820-w

81 Hippen, A. A., et al. miQC: An adaptive probabilistic framework for quality control of single-cell RNA-sequencing data. PLoS Comput Biol 17, e1009290 (2021). 10.1371/journal.pcbi.1009290

